# SMSTPred: Protein Language Model Embeddings Predict SAM-Methyltransferase Substrate Specificity and Reveal Metabolite Methyltransferasome Enrichment in Plants and Actinomycetota

**DOI:** 10.64898/2026.11.13.766537

**Authors:** Nathaniel Roethler, Dmitrij Rappoport, Ricardo Almada Monter, Adriana Siordia, Javier Espinoza Herrera, Adrian Jinich

## Abstract

S-adenosyl-methionine (SAM)-dependent methyltransferases (MTases) are one of the largest and most ancient enzyme superfamilies in biology, transferring a methyl group from SAM to substrates as different as DNA, RNA, proteins, and small-molecule metabolites. Most SAM-MTases share a Class I Rossmann-like SAM-binding core, with substrate specificity determined by variable inserts and accessory domains rather than the conserved fold; as a result, substrate type is unknown for most MTases in sequence databases, which are dominated by unreviewed entries. Here we present SMSTPred (SAM-Methyltransferase Substrate-Type Predictor), a multi-label classifier that predicts MTase substrate type from protein language model embeddings and benchmarks it under homology-disjoint cross-validation against sequence-homology, structure-homology, and profile-HMM baselines. SMSTPred reaches a cluster-disjoint Matthews correlation coefficient (MCC) of 0.94, significantly ahead of sequence-homology (MMseqs2, 0.85), structure-homology (Foldseek, 0.89) and profile-HMM (0.52) baselines, while providing calibrated, per-substrate confidence. Applying it to ∼116,000 MTases across 1,249 reference proteomes, we map the substrate landscape of methylation across the tree of life. Substrate specialization tracks taxonomy: animals expand protein- and DNA-methylation, whereas small-molecule methyltransferases are enriched in plants and Actinomycetota; in Actinomycetota beyond what is expected for their proteome size, and in plants in proportion to their larger proteomes. The predictions identify thousands of uncharacterized methyltransferases with substrate type assignments (about half of those in uncharacterized clusters), many corroborated by an independent reaction-retrieval method. Notably, ∼11% of enzymes with a single substrate call lack any class-discriminative sequence motif, suggesting that substrate specificity is often encoded beyond short linear motifs. Together, this work provides SMSTPred, a substrate-type predictor for Class I methyltransferases, and a tree-of-life resource of testable functional hypotheses.

## Introduction

Methylation is among the most widespread modifications in biology, and most of it is carried out by SAM-dependent methyltransferases using SAM as the methyl donor.^1–3^ Methyltransferases serve a variety of biological roles: DNA methylation drives epigenetic regulation and bacterial restriction-modification;^4–6^ RNA methylation underlies the epitranscriptome and the maturation of ribosomal, transfer, and messenger RNAs;^7–9^ protein methylation shapes chromatin through histone and arginine marks and tunes signaling and translation;^10–13^ and small-molecule methylation builds and tailors secondary metabolites, natural products, hormones, and cofactors.^14^ The same catalytic chemistry therefore participates in completely different pathways depending on the substrate, making substrate type a key functional characterization of the superfamily.^15,16^

The substrate of the vast majority of MTases in sequence databases is unknown.^17^ Of the ∼1.6 million UniProtKB proteins assigned to the Class I SAM-MTase superfamily (InterPro IPR029063), fewer than 0.7% are manually reviewed, and only ∼1,300 (0.08%) carry a catalytic activity supported by experimental evidence. Prediction tools are therefore needed, but predicting substrate types at scale is hard. Most SAM-MTases share a Class I Rossmann-like core whose conserved motifs are constrained by binding the common cofactor, SAM, rather than the substrate.^3,18^ The methyl-acceptor pocket is shaped instead by variable loops and inserts at the edge of this core and by appended domains, such as RNA- and DNA-recognition modules, that differ widely between families.^3,14,18^ As a result, homologous Class I cores can methylate chemically unrelated substrates, from small molecules to nucleic acids and proteins, while the same substrate class can be methylated by structurally unrelated folds, as with protein lysine methylation by both SET-domain and Class I enzymes.^3,13^ Global sequence or structural similarity is therefore an unreliable guide to substrate, and homology-based transfer is limited to enzymes with a characterized relative, which, given the numbers above, excludes much of the superfamily.

Existing computational tools fall short for this reason. Detecting the Class I SAM-MTase domain, or a remote homolog, does not by itself establish substrate type: closely related Class I enzymes can act on different substrate classes, and homology- or profile-based substrate assignment is correspondingly incomplete.^19–25^ The release-factor methyltransferase HemK, for instance, carries a DNA-amino-methyltransferase signature yet methylates protein.^26^ Prior substrate-type classifiers were built for a single organism’s methyltransferasome from hand-crafted features and evaluated in-sample or by leave-one-out that was not homology-controlled, so their reported accuracies do not establish how well they generalize to enzymes without a characterized relative — the regime that covers almost the entire superfamily.^22,23,27,28^

Protein language models (PLMs) offer a different route to this problem. Trained by self-supervision on hundreds of millions of sequences, they map each protein to a fixed-length vector (an embedding) without alignments or experimentally determined structures.^29–31^ These embeddings carry information on secondary structure, residue contacts and function, and they allow annotation transfer between proteins whose sequences are too dissimilar for alignment-based methods.^29,32,33^ Models that also encode structure-derived information extend this further.^34–36^ For enzymes, supervised classifiers on PLM representations predict Enzyme Commission numbers more accurately than BLAST-based homology transfer, especially for sequences with distant relatives.^37^ These results suggest, but do not establish, that PLM embeddings could capture substrate-relevant features in the variable regions of Class I MTases that sequence and structural homology tools miss.

To test this hypothesis, we built a multi-label substrate-type classifier (DNA, RNA, Protein, Metabolite substrates, metabolite being any biological compound that does not fall into DNA, RNA, or Protein) on protein language model embeddings. We benchmark this classifier with homology-disjoint cross-validation against sequence-homology, structural-homology, and profile-HMM baselines. We deploy it across 1,249 reference proteomes and roughly 116,000 methyltransferases to map the evolutionary landscape of methylation substrate specialization, and we show that it recovers canonical enzyme families de novo while surfacing thousands of uncharacterized, substrate-type predicted enzymes, a resource for discovery and annotation (Figure 1). We also release the trained model as SMSTPred, a ready-to-run classifier for the substrate type of any methyltransferase sequence.

**Figure 1.**
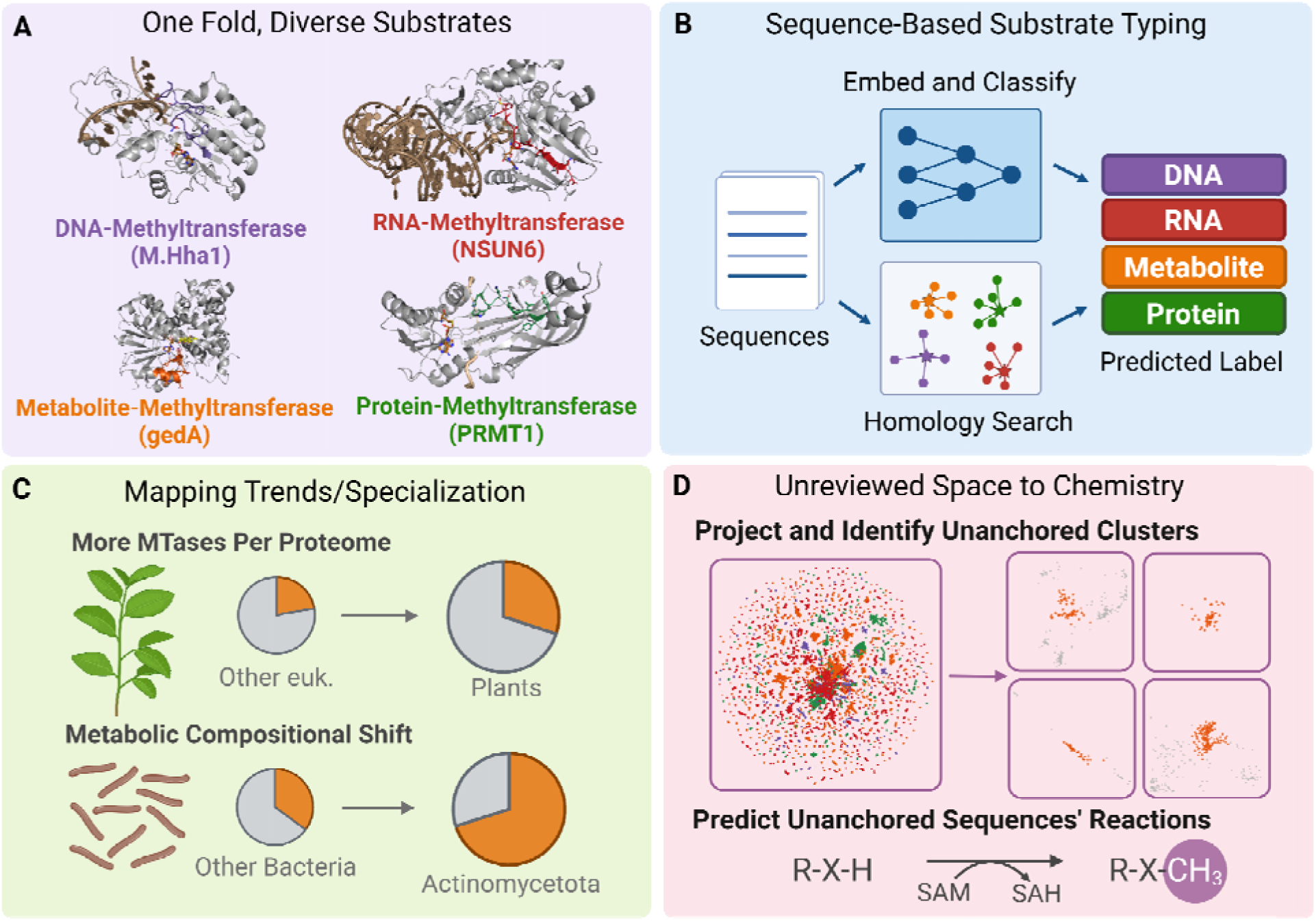
(A) Class I SAM-dependent methyltransferases share a Rossmann-like fold and act on DNA, RNA, proteins, or small-molecule metabolites; the structures show enzymes bound to a substrate of each class. (B) Two routes from sequence to substrate type. A multi-label classifier takes the ProtT5 embedding of a sequence and returns an independent probability for each of the DNA, RNA, Metabolite, and Protein classes (cluster-disjoint cross-validation, MCC 0.94). Homology-based assignment instead transfers the label of the closest annotated enzyme identified by sequence search (MMseqs2, profile HMM) or structure search (Foldseek), and makes no call when no sufficiently similar annotated enzyme exists. (C) The classifier was applied to 115,850 SAM-MTase sequences from 1,249 reference proteomes. The proportion of enzymes predicted to act on metabolites is higher in plants than in other eukaryotes (median 42% versus 34%) and higher in Actinomycetota than in other bacteria (72% versus 35%). (D) UMAP projection of the ProtT5 embeddings, colored by predicted substrate type. Clusters without any reviewed (Swiss-Prot) member were assigned an Enzyme Commission number with CLEAN and a best-scoring Rhea reaction with Horizyn; in these reactions SAM donates a methyl group to the substrate and is converted to S-adenosylhomocysteine (SAH).

## Methods

SMSTPred denotes the deployed model: a six-member ensemble of multi-label feed-forward networks on mean-pooled ProtT5 embeddings, classified at a fixed probability of 0.5 for every label, with an ensemble-disagreement confidence and an applicability-domain flag (Deployment model and confidence).^29,38–40^ MLP_ProtT5 denotes the same architecture when evaluated in cross-validation; the MLPs on the other embeddings and the homology-based baselines were benchmarked for comparison and are not part of SMSTPred.

### MTase Training Set Curation

We used a four-way, multi-label scheme (DNA, RNA, Protein, and Metabolite substrates). Reviewed UniProtKB/Swiss-Prot entries (release 2026_03) annotated as SAM-dependent methyltransferases (IPR029063) were retrieved; entries with several catalytic-activity annotations were collapsed to one record carrying the union of their substrate labels, giving 7,553 unique proteins of up to 1,000 aa (7,582 of up to 2,000 aa for the deployed model). Substrate assignments were taken from curated catalytic-activity annotations (UniProt reaction text with Rhea and ChEBI identifiers) for each SAM-dependent reaction supported by a literature reference or a curated rule (HAMAP, ProRule, RuleBase).^17,41^ Enzymes acting on more than one substrate were handled as multi-label. The final training set is strongly imbalanced (RNA-rich, DNA-rare: 5,017 RNA, 1,419 Metabolite, 1,014 Protein, and 110 DNA positive labels).

### UniProt (UP) Training Set Curation

For the training-set-scale comparison (Figure 2B), we additionally assembled a broad comparison set of 204,941 reviewed (Swiss-Prot; release 2026_03) enzymes of up to 1,000 aa under the same four-way substrate scheme, not restricted to SAM-methyltransferases (UP).^17,42^ Labels were assigned by the same catalytic-activity annotation pipeline used for the focused set (MTase), and all 7,553 SAM-MTases of the focused set are themselves members of UP, with identical labels. UP is about 27-fold larger but far more label-imbalanced than the focused set (approximately 142,000 Metabolite, 36,000 RNA, 26,000 Protein, and 4,200 DNA positive labels), and is used solely to test whether breadth of training data is more effective than task-matched specificity.

**Figure 2.**
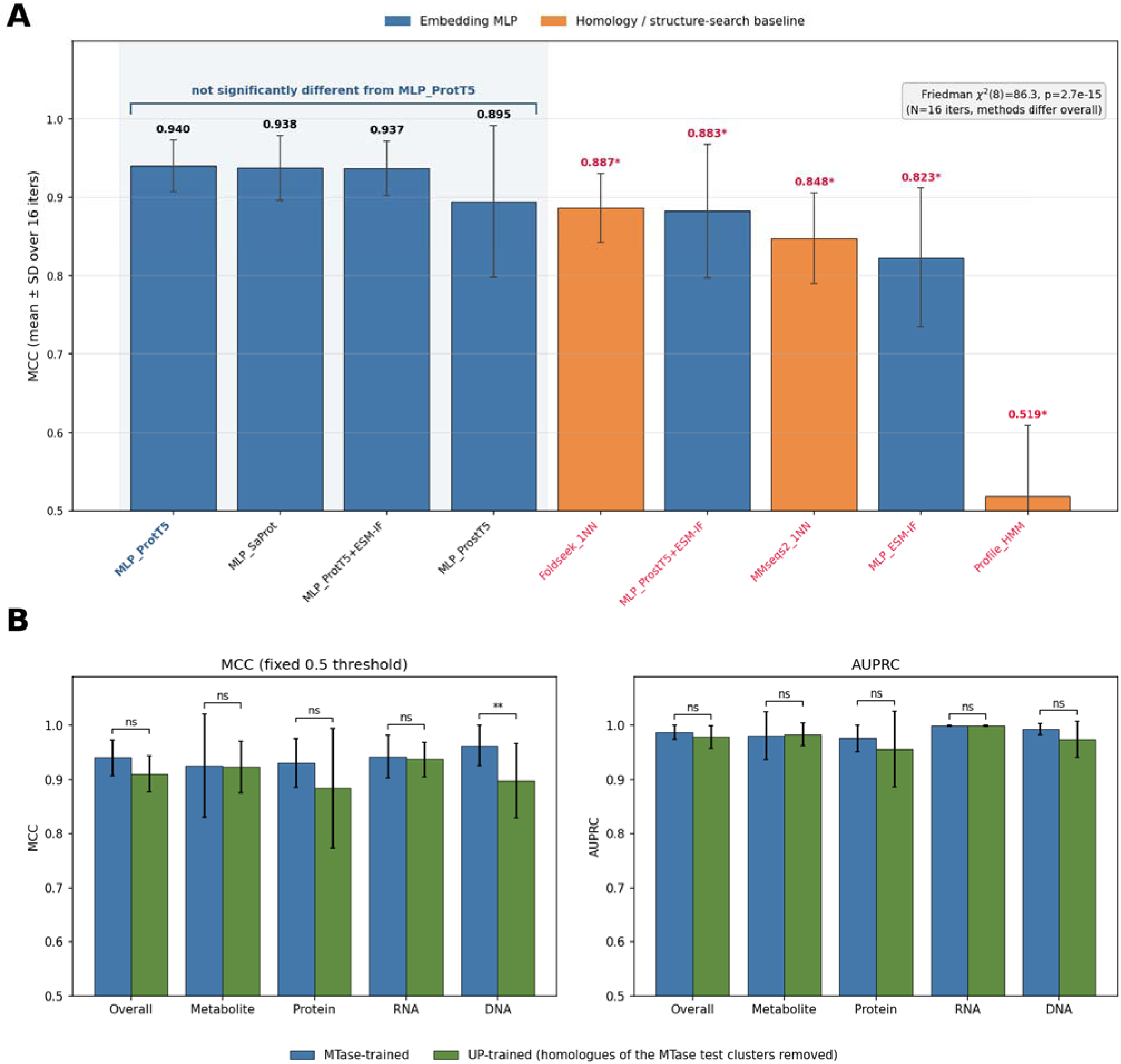
Benchmarking of the classifier. (A) Cluster-disjoint MCC for the embedding MLPs and the homology-search baselines (MMseqs2 and Foldseek 1-nearest-neighbor, and a profile HMM), mean +/- SD over sixteen cluster-disjoint splits, each scored on the same held-out sequences. MCC is averaged over the four substrate labels at a fixed threshold of 0.5. A Friedman omnibus test (chi-square(8) = 86.3, p = 2.7 x 10^-15^) was followed by paired Wilcoxon signed-rank tests against the deployed MLP_ProtT5 with Holm correction; red values marked with a star are significantly worse than MLP_ProtT5. (B) MTase-trained (blue) versus UP-trained (green) ProtT5 MLP, scored on held-out

### Tree-of-life prediction set curation

The prediction set spanned reference proteomes for 1,249 organisms across bacteria, archaea, and eukaryotes, with taxonomy metadata (UniProtKB (Swiss-Prot and TrEMBL); release 2026_03).^17,42^ Candidate methyltransferases were identified by InterPro SAM-binding domain membership plus a length filter (up to 2,000 aa), yielding 115,850 candidate methyltransferase sequences (a further 1,703 domain-bearing proteins longer than 2,000 aa were not scored).^17^

### Protein Language Models employed

We compared a sequence protein language model (ProtT5), a structure-aware sequence model (ProstT5, which is informed by 3Di structure tokens), a structural / inverse-folding model (ESM-IF), and a structure-token model (SaProt), along with concatenations.^29,34,35,43^ Each protein is represented by the mean of the model’s per-residue embeddings (ProtT5 and ProstT5, 1,024 dimensions; ESM-IF, 512; SaProt, 1,280). ProtT5 (Rostlab/prot_t5_xl_half_uniref50-enc, the half-precision encoder) and ProstT5 (Rostlab/ProstT5 encoder, <AA2fold> prefix) need only the amino-acid sequence at inference, whereas ESM-IF (esm_if1_gvp4_t16_142M_UR50) and SaProt (SaProt_650M_AF2), and any concatenation containing ESM-IF, also need a structure. The deployed model uses the ProtT5 embedding, which only requires sequence.

### Structures curation

Structures were drawn from the AlphaFold Database, supplemented by structures predicted locally with ColabFold 1.6.1 (AlphaFold2-ptm, five models, MSAs from the ColabFold MMseqs2 server, no templates or relaxation; the model with the highest pLDDT was used) where AlphaFold DB had no current model (in release 2026_03: 143 AlphaFold DB models fetched anew, 9 ColabFold predictions, and 7,401 structures reused from earlier releases after their sequence was checked).^44–46^ Structures were needed only for the ESM-IF/SaProt benchmark arms; deployment uses ProtT5 from sequence alone.

### Classifier architecture and training

SMSTPred is built on a multi-label feed-forward network on the embedding, predicting independent per-substrate probabilities, implemented in PyTorch.^47^ Binary cross-entropy on the sigmoid of each logit was used, so the four labels were predicted independently. The MLP maps the embedding through two fully connected hidden layers (512 and 256 units, ReLU activation, no dropout or weight decay) to four logits, one per substrate class (DNA, RNA, protein, metabolite), and was trained with Adam (learning rate 1e-3, one gradient step per epoch on a random 90% of the training proteins). Learning rate, network size and patience were fixed in advance and not tuned. Training stopped early when the validation loss on held-out clusters had not improved for 80 epochs (at most 2,000 epochs), and the epoch with the lowest validation loss was kept (median 204 epochs for ProtT5; Fig S1); the UP-trained model was trained in the same way (Fig S2). The same architecture, with the input width set by the embedding, was used for every embedding in the benchmark.

### Homology-aware evaluation

The training set was clustered with MMseqs2 (easy-cluster; 30% sequence identity, alignments covering at least 80% of both sequences, most sensitive setting, -s 7.5), which gave 502 clusters for the 7,553 benchmark proteins (the largest holds 830 proteins; 248 are singletons).^48^ Cross-validation was cluster-disjoint, so that training and test folds never share a homology cluster: in each of sixteen repeats whole clusters were assigned at random, 20% to testing, 15% to validation and the rest to training (test sets of 1,200 to 2,397 proteins, median 1,634; training sets of 4,345 to 5,775). MCC was computed at a fixed decision threshold of 0.5 for every label.^49,50^

### Benchmarking design

Performance was estimated on sixteen random partitions of the sequence clusters into training (65%), validation (15%) and test (20%) sets, so that no test protein shares a cluster with a training or validation protein. Epochs were selected by early stopping on the validation clusters only; the test clusters played no role in training or model selection. All methods were trained and evaluated on the same partitions, so their test scores were paired within each partition and were compared with paired tests. The classifier comparison scored the embedding MLP against a profile-HMM, a sequence-homology nearest-neighbor (MMseqs2), and a structural-homology nearest-neighbor (Foldseek).^48,51,52^ All methods were scored on identical held-out protein clusters, over sixteen repeated iterations, with MCC as the primary metric alongside AUPRC, AUROC, F1, and accuracy.^49,53–56^ We ran a Friedman test across the sixteen splits on the methods of Figure 2A and compared each method with the deployed MLP_ProtT5 using a paired Wilcoxon signed-rank test on the sixteen MCC differences, with a Holm correction across the eight comparisons of Figure 2A.^57–59^ Because the sixteen iterations share training data, these rank tests could slightly overstate the significance of the differences. The MTase-versus-UP comparison used paired Wilcoxon signed-rank tests over the sixteen splits (Holm-corrected across the five comparisons) and, as a stricter check that corrects for the overlap between training sets, a Nadeau-Bengio corrected resampled paired t-test.^60,61^

### Deployment model and confidence

The deployed model (SMSTPred) is an ensemble of six neural networks with the same architecture (the multi-label MLP described above), each trained on a different part of the 7,582 deployment proteins. To build it, the sequence clusters were divided into six folds of similar size. Each network is trained on five folds and validated on the sixth, a different fold for each member. The models were trained using early stopping, as described above. All six trained models were deployed on sequences we were predicting on, with the mean of the six members’ predicted probabilities used for classification. The standard deviation of the six probabilities (their disagreement) also served as an uncertainty estimate. Labels were assigned at a fixed decision threshold of 0.5. Out-of-fold performance was estimated with separate audit members whose test fold was held out from training and epoch selection (see Results). Each protein receives per-substrate scores, multi-label calls, an ensemble-uncertainty, and an applicability-domain flag that marks a protein as out of domain when its highest cosine similarity to any training protein in ProtT5 space falls below 0.81 (the 1st percentile of the training proteins’ own nearest-neighbor similarities).

### Tree of Life and UMAP cluster analysis

The predictions for the 1,249 reference proteomes were analyzed in two ways. To look for taxonomic trends, we built a taxonomic tree of the proteomes from the NCBI lineage of each organism (unit branch lengths, so the tree shows taxonomic relationships rather than sequence phylogeny). Each leaf was annotated with the fraction of that organism’s predicted SAM-MTases called DNA, RNA, protein, or metabolite (a protein with two calls counts toward both), the number of SAM-MTases, and their share of the proteome. The tree was drawn in a circular layout with one ring per substrate class, so that lineages enriched in a class appear as contiguous sectors. The per-organism compositions were compared between taxonomic groups with Mann-Whitney U tests (two groups) or Kruskal-Wallis tests (more than two groups).^62,63^ To relate the predictions to protein families, we projected the ProtT5 embeddings of all predicted SAM-MTases into two dimensions with UMAP (15 neighbors, minimum distance 0.1, cosine metric), so that proteins with similar embeddings lie close together, and grouped the projection into density-based clusters with HDBSCAN (minimum cluster size 50, minimum samples 10) applied to the two-dimensional coordinates.^64,65^ Because UMAP and HDBSCAN are stochastic, the map was computed once and kept fixed for all downstream analyses. For each cluster we tabulated the predicted substrate classes and taxa, and the reviewed (Swiss-Prot) members that tie the cluster to characterized enzymes; clusters without any reviewed member were treated as uncharacterized.

To validate the substrate classes against known protein families, InterPro domain enrichment was computed per class over the 114,861 proteins with a single class label: every InterPro domain present in at least 20 proteins of a class was tested against all other classes pooled with a one-sided Fisher’s exact test on domain presence versus absence, and domains with at least two-fold higher prevalence in-class than in the pooled background were retained and ranked by in-class count (Table S3).^66^ Domains were selected by fold enrichment and prevalence, and the Fisher p-values were not corrected for multiple testing.

To test whether lineage-level substrate specialization simply tracks proteome size, we additionally fitted power laws n = λ g, with n the number of predicted SAM-MTases (total, or with a Metabolite call) and g the proteome size, separately for Actinomycetota and the other bacteria and for each of four eukaryotic clades (Viridiplantae, Fungi, Metazoa, and the remaining eukaryotes, assigned from the NCBI lineage of each proteome; archaea were too few to fit). Proteomes were grouped into logarithmic size bins (0.15 decades), and bins with fewer than five proteomes were dropped. The protocol of De Lazzari et al. (2017) requires at least ten genomes per bin, which leaves a single bin for Viridiplantae and prevents a fit; a minimum of five proteomes per bin was therefore used for all clades. The exponent α and the prefactor λ were estimated by non-linear least squares weighted by the standard error of each bin mean.^67^ The 95% intervals of the exponents and of the fitted counts come from 1,000 bootstrap replicates that resample whole clades (lineage prefixes of depth seven in bacteria and eight in eukaryotes) rather than proteomes, to allow for phylogenetic non-independence; a phylogenetic regression on a dated tree was not performed. To test whether clades differ in the number of SAM-MTases at a given proteome size or in how that number scales with proteome size, a single power law for all proteomes of a domain was compared with clade-specific power laws. The base-10 logarithm of the count was regressed on the logarithm of proteome size by least squares with (i) one line for all proteomes of the domain, (ii) clade-specific intercepts and a common slope, and (iii) clade-specific intercepts and slopes; the models were compared by R² and the Akaike information criterion. In contrast to De Lazzari et al., who compare the exponent of each family with that of its functional category, each clade was fitted separately and clades were compared with one another, because a single power law fitted to all non-plant eukaryotes fitted poorly (reduced χ² of 29 for total and 54 for Metabolite counts) and clades differed in level. Each clade was fitted by weighted non-linear least squares, with goodness of fit assessed by the reduced χ², and the fitted counts of the clades were compared at a reference proteome size within the size range of every clade of the domain (6,000 proteins in bacteria, 20,000 in eukaryotes). The exponents describe the clades over the observed range of proteome sizes and are not general scaling laws.

### CLEAN and Horizyn analysis

In this study we use the term ‘dark clusters’ to refer to HDBSCAN clusters of the ProtT5 UMAP that contain no reviewed (Swiss-Prot) member and in which at least 80 percent of the proteins are both from a single lineage (Actinomycetota or plants) and predicted ‘Metabolite substrates’. Of 947 clusters, 559 have no reviewed member, and 53 of these meet this criterion (23 Actinomycetota, 30 plant; 5,216 proteins). For the uncharacterized (’dark’) clusters with no reviewed anchor, we cross-validated the SMSTPred substrate calls with two independent, pretrained methods that use neither our classifier nor our labels. CLEAN assigns each dark protein the EC number of its nearest enzyme centroid in a contrastively-trained embedding space and reports the distance d from the protein to that centroid; the call of a dark protein cluster is the EC number assigned to most of its members, with a confidence tier from the median d of its members (reliable below 6, marginal 6 to 8, unreliable above 8). The EC number was named from the ExPASy ENZYME database to judge whether it names a genuine methyltransferase, and annotated with the UniProt pathway of the characterized enzymes that carry that EC.^37^ Horizyn, a pretrained dual-encoder contrastive reaction-retrieval model, embeds a reaction (RDKit+/DRFP fingerprints) and a protein (ProtT5-XL) into a shared space and ranks candidate reactions by cosine similarity.^68^ We screened every dark-cluster protein against all 36,552 scored Rhea reactions and, because Rhea encodes one chemistry as many near-duplicate reactions, additionally clustered all scored reactions in Horizyn’s own embedding space (UMAP + HDBSCAN, as above) into reaction families, assigning each protein cluster the family that contains the best-scoring reaction of most of its members. The reaction assigned to a protein cluster is the best-scoring reaction shared by most of its members; this per-cluster assignment is independent of the ensemble used for substrate-type prediction. The clusters shown in Figure 8 were chosen by rule (CLEAN reliable, a methyltransferase EC, and that EC inside Horizyn’s dominant reaction family).

### Conserved per-label motif analysis

To interpret conserved motifs from SAM MTase predicted labels, we first assembled a labeled corpus of 122,807 proteins (the 7,553 benchmark proteins with curated labels plus the tree-of-life proteins labeled by SMSTPred), kept the 114,861 proteins with exactly one substrate class, and reduced them to family representatives by 50 percent-identity clustering (MMseqs2). For each substrate class, we used STREME to find short motifs enriched in that class versus the other three (eight motifs per class, thirty-two total).^69^ Because the substrate-type labels are the classifier’s own predictions, the motifs describe what SMSTPred assigns; only the training proteins carry curated labels. We then scanned every motif with FIMO at p < 1e-4 against all 122,807 proteins to obtain the real per-protein coverage of each motif and to compute its class specificity, defined as the occurrence rate in its own class divided by the occurrence rate in all other proteins pooled, where the occurrence rate of a motif in a group of proteins is the fraction of proteins in the group with at least one match.^70^ The class-discriminative set was defined by class specificity, not by the STREME E-value, which tests enrichment against the control sequences at discovery and does not measure how often a motif occurs outside its class. Motifs with a specificity of at least two (twenty-four) were retained, and the remaining eight were classified as pan-methyltransferase, Rossmann-core signatures shared across classes. Finally, for each class-specific motif we searched every FIMO carrier that has an experimental structure (UniProt and RCSB records) for entries holding both a SAM-like cofactor and a substrate, tried them in a fixed order (own class, reviewed, FIMO p-value, resolution), located the motif by its FIMO matched sequence, measured its minimum heavy-atom distance to the cofactor and to the substrate, and rejected structures in which a ligand sits on another chain or away from the active site. Of 366 motif–structure pairs evaluated, 181 (96 distinct PDB entries) passed these criteria, with one to eight passing structures per motif (the search stopped at eight). The resulting structures were ranked by rule (a substrate of the class’s own kind, own class, the placement most of the motif’s structures show, reviewed, intact SAM, resolution), and the first was used to classify the placement of the motif (near both, near SAM, near substrate, or distal; contact is 4 Å or less).

### Software, data, and availability

SMSTPred code, trained model weights and per-organism predictions: https://github.com/jinichlab/SMSTPred (archived at https://doi.org/10.5281/zenodo.24186207). Embeddings, label tables, structures and tree-of-life resources: Zenodo, https://doi.org/10.5281/zenodo.23655998. A ready-to-run version of the classifier, with a walkthrough for preparing input sequences and reading the output, is in the ready_classifier folder of the repository and always uses the newest trained model. The entire analysis, UniProt pull to every figure and table, is one automated pipeline (workflow/run_refresh.sh) that can be rerun for each UniProt release.

## Results

### Protein language model embeddings predict methyltransferase substrate class with high accuracy

To determine whether protein language model embeddings can accurately distinguish methyltransferase substrate classes without relying on explicit 3D structures, we systematically benchmarked classifier heads across embeddings under homology-disjoint evaluation. All heads have the architecture described in the Methods. The comparison of the other heads with the deployed ProtT5 head is reported below. No embedding led on every label: ProtT5 was best on DNA (0.963), Metabolite (0.926) and Protein (0.931), and SaProt on RNA (0.952). Concatenating embeddings added little (Figure 2A). Training stopped early at a median of 204 epochs for ProtT5 (range 154 to 596; Figure S1), and the rare DNA head remained the principal source of variance.

MTase proteins over the same sixteen splits (mean +/- SD), per substrate class and overall (mean of the four classes), with Wilcoxon signed-rank significance (Holm-corrected across five comparisons). Only DNA differs after correction, and it does not under the stricter Nadeau-Bengio corrected test.

Under cluster-disjoint evaluation, the deployed ProtT5 MLP reached a mean MCC of 0.940 over the four labels (Figure 2A). The nine methods of Figure 2A differed significantly overall (Friedman chi-square(8) = 86.3, p = 2.7 x 10^-15^). In paired Wilcoxon signed-rank tests against ProtT5 with Holm correction, SaProt (0.938), ProtT5+ESM-IF (0.937) and ProstT5 (0.895; SD across splits 0.097) were not significantly different (Holm p = 0.46, 0.39 and 0.22), whereas Foldseek (0.887; Holm p = 0.002), the ProstT5+ESM-IF concatenation (0.883; p = 0.007), sequence-homology nearest neighbor (MMseqs2, 0.848), ESM-IF alone (0.823) and the profile HMM (0.519) were significantly worse (the last three p = 2.4 x 10^-4^). We deploy ProtT5, which needs only the sequence rather than a structure as input. Homology search remains a serious but clearly weaker competitor: MMseqs2 and Foldseek trail the deployed model by 0.05 to 0.09 MCC. Sequence-derived representations matched or outperformed explicit-structure ones (ProtT5 0.940 vs ESM-IF 0.823), and adding ESM-IF to ProtT5 gave no gain. This indicates that substrate specificity lives more in sequence-level recognition determinants than in the conserved global fold.

Finally, we asked whether broader training data would increase accuracy. A larger set could add sequence diversity and data points, but its labels mostly reflect other enzyme families, so it could equally dilute the methyltransferase-specific signal. The UP set contains all 7,553 methyltransferases plus about 197,000 other proteins, with homologues of the test clusters removed. Training the same ProtT5 MLP on the MTase set (MTase-trained) or on the 204,941-protein UP set (UP-trained) and evaluating both on the same held-out MTase proteins gave overall MCCs of 0.940 and 0.911 (Wilcoxon p = 0.016, Holm p = 0.062; Nadeau-Bengio p = 0.38). Only the rare DNA class differed after correction (0.963 vs 0.898; Holm p = 0.007), and not under the stricter Nadeau-Bengio test (p = 0.15); Metabolite (0.926 vs 0.923), Protein (0.931 vs 0.884) and RNA (0.942 vs 0.937) did not differ (Figure 2B). The additional enzymes therefore did not improve accuracy.

We deployed SMSTPred, the six-member ProtT5 ensemble (Methods), on the 115,850 candidate methyltransferases of the 1,249 reference proteomes. Out-of-fold performance of separate audit members on the deployment proteins was MCC 0.955 (Metabolite 0.962, Protein 0.913, RNA 0.969, DNA 0.976). The threshold was 0.5 for every label, so no per-label tuning is needed; this estimate is not directly comparable to the cluster-disjoint MCC of 0.940 in Figure 2A. Each prediction carries the ensemble standard deviation as an uncertainty and an applicability-domain flag based on the cosine similarity to SMSTPred’s training set (cutoff 0.81); 43.7% of the proteins fall outside the domain, and their calls are of lower confidence. The calls are analyzed across the tree of life below.

### Substrate specialization across the tree of life tracks taxonomy

Gene counts in functional categories are known to scale with genome size in category-specific ways; here we can ask the same question within a single fold family, resolved by substrate.^67,71–73^ Applying SMSTPred to 115,850 enzymes across 1,249 reference proteomes produced a global substrate-class map (Figure 3). Small-molecule (Metabolite) methyltransferases are the most abundant class (about 47,400 enzymes, 40.9% of the total), followed by RNA (about 36,700; 31.7%), Protein (about 20,700; 17.9%), and DNA (about 7,300; 6.3%). Of all proteins, 93.1% received a single class call, 1.8% received more than one, and 5.0% fell below the threshold for every class and are reported as unassigned. This inverts the composition of the reviewed Swiss-Prot set, which is RNA-biased, suggesting that the RNA bias reflects which enzymes have been studied and curated rather than their natural abundance.

**Figure 3.**
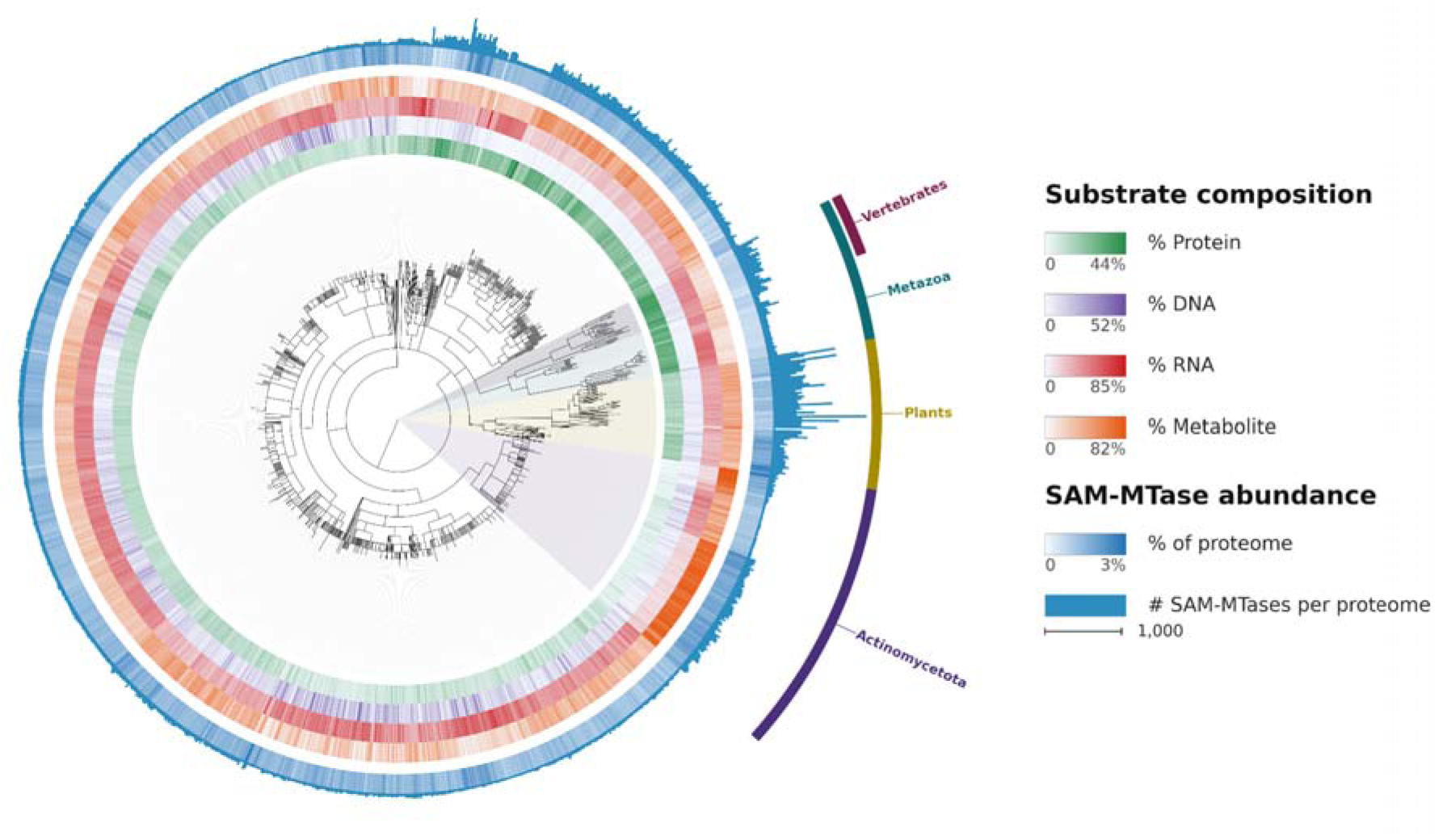
Tree-of-life landscape. Substrate composition of the predicted SAM-methyltransferases of 1,249 reference proteomes, arranged on the NCBI taxonomy (circular tree with unit branch lengths, showing taxonomic relationships). Rings, inner to outer: percentage of each proteome’s SAM-methyltransferases predicted as Protein, DNA, RNA and Metabolite (a protein with two calls counts toward both) and, after a gap, the percentage of the proteome that is SAM-methyltransferase. Outer bars: number of SAM-methyltransferases per proteome (scale bar, 1,000). Color scales run from 0 to the maximum given in the legend.

Metazoa are enriched in protein-methyltransferases (PRMT and CARM1 families) relative to other eukaryotes, both by count (median 51 per proteome versus 24 in unicellular eukaryotes and 38 in non-metazoan multicellular eukaryotes) and by proteome fraction (36.7% vs 23.1% and 24.0%; Figure 4A).^12^ This result is consistent with the proposal that expanded protein modification and regulation accompanied the evolution of eukaryotic multicellularity.^74^ Within Metazoa, vertebrates show a marked expansion of DNA methyltransferases relative to invertebrates, both by count (median 19 vs. 2; Mann-Whitney U = 886.5, p = 2.8e-10) and by proteome fraction (6.9% vs. 2.2%; U = 874, p = 1.0e-9), alongside elevated protein methylation by count (median 95 vs. 37; U = 864, p = 2.5e-9) though not by proteome fraction (37.0% vs. 36.5%; U = 479, p = 0.73) (Figure 4B). This supports previous conclusions that DNA and Protein methylation are essential in the development of neurons and a central nervous system.^75–77^

**Figure 4.**
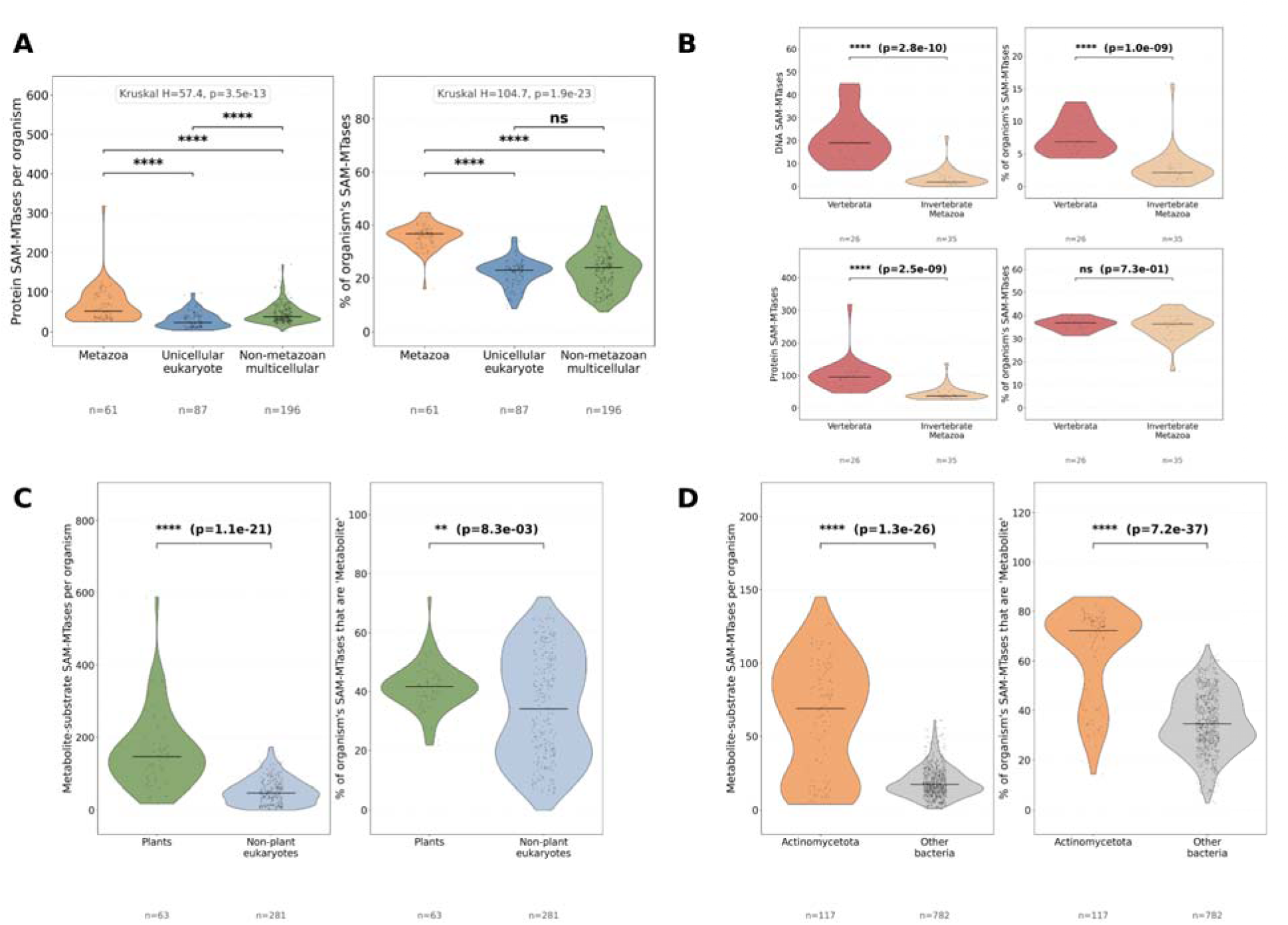
(A) Protein-methyltransferase differentiation across metazoa, unicellular, and non-metazoan multicellular eukaryotes; (B) vertebrate versus invertebrate DNA and protein methyltransferase differentiation; (C) elevated small-molecule (Metabolite) methyltransferasome in plants versus non-plant eukaryotes; (D) the same in Actinomycetota versus other bacteria. Each panel shows per-proteome counts and percentages with Mann-Whitney U tests.

Plants and Actinomycetota both carry strongly elevated ‘Metabolite substrates’ (small-molecule) methyltransferase counts, relative to other eukaryotes and other bacteria respectively (plants p = 1.1e-21; Actinomycetota p = 1.3e-26, both by Metabolite count; Figure 4C,D). The counts were related to proteome size by fitting the number of predicted SAM-methyltransferases as a power law of proteome size separately for each clade (Figure 5). In bacteria, the total count of Actinomycetota increases with a higher exponent than that of other bacteria (1.42 [0.99, 1.64] versus 0.69 [0.60, 0.83]), and the fitted count of Metabolite-substrate enzymes at 6,000 proteins is 53 [45, 69] in Actinomycetota and 28 [26, 35] in other bacteria (Figure 5A,B). In eukaryotes, the fitted count of Metabolite-substrate enzymes at 20,000 proteins is 105 [90, 170] in plants, 150 [69, 192] in fungi, 50 [26, 86] in the remaining eukaryotes, and 12 in metazoa (Figure 5D). The intervals of plants and fungi overlap, and both lie above that of metazoa. The exponents of the eukaryotic clades are poorly constrained, and the fits differ in quality: the reduced χ² is 0.9 to 1.5 for plants and 14 to 20 for fungi (Figure 5C,D), and a single power law fitted to all non-plant eukaryotes fits poorly (reduced χ² 29 for total and 54 for Metabolite counts). Clades were therefore fitted separately: a separate intercept for each clade increased R² for Metabolite counts across eukaryotes from 0.28 (one line for all eukaryotes) to 0.74, and clade-specific slopes added little (0.76). Small-molecule methyltransferases are thus more numerous at equal proteome size in plants and fungi than in metazoa, and in Actinomycetota than in other bacteria; these data do not distinguish plants from fungi. O-methyltransferase and SABATH domains are among those enriched in the predicted ‘Metabolite substrates’ class overall (Table S3).

**Figure 5.**
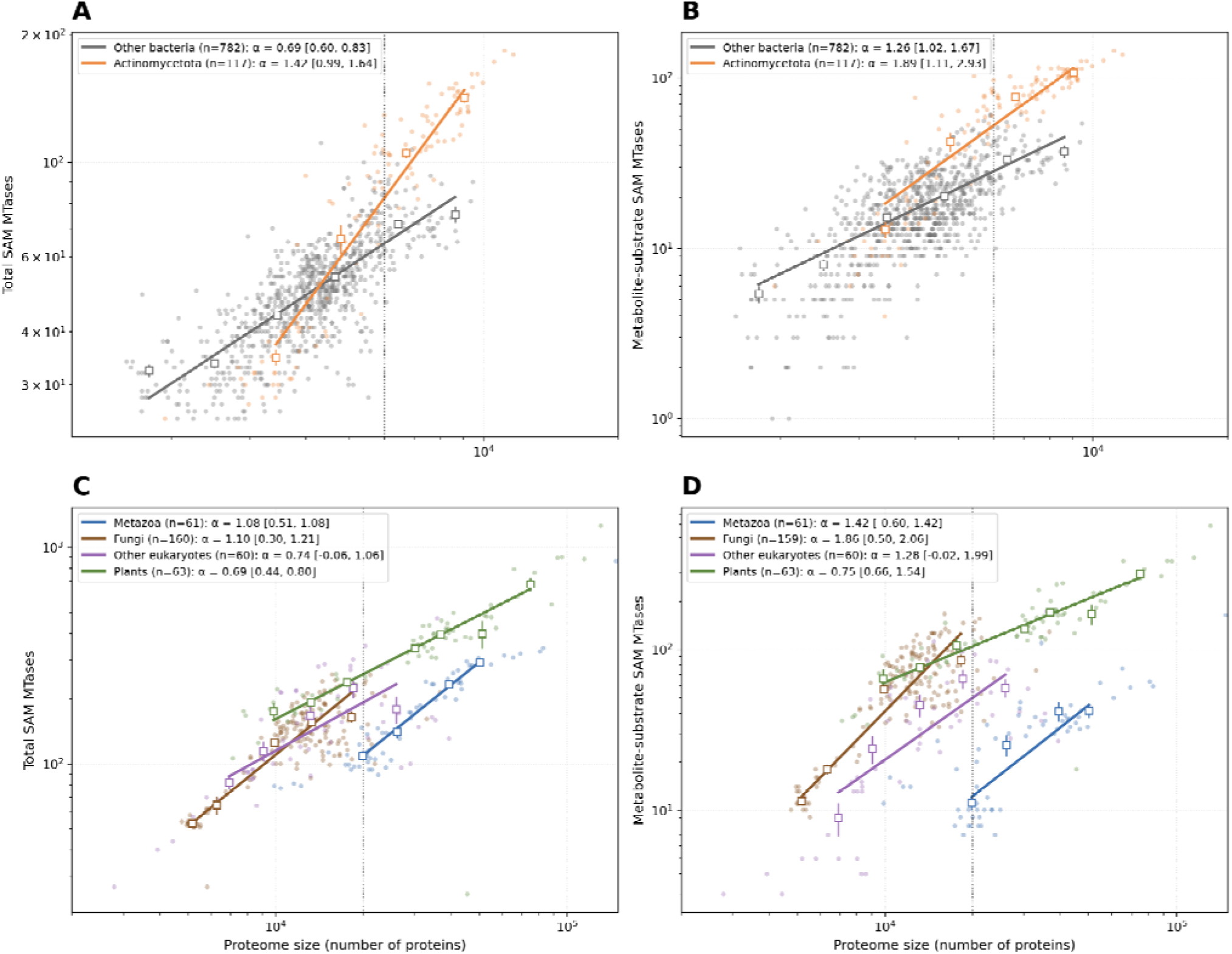
Proteome size and SAM-methyltransferase counts, fitted separately for each clade as n = λ g (n, number of predicted SAM-methyltransferases; g, proteome size). (A, B) Actinomycetota and other bacteria; (C, D) plants, fungi, metazoa, and other eukaryotes. Panels A and C show all predicted SAM-methyltransferases and panels B and D those with a Metabolite call. Small markers are proteomes, open markers the mean (± s.e.) of logarithmic size bins (0.15 decades, at least five proteomes), and lines weighted non-linear least-squares fits. The legends give the exponent α of each clade with its 95% interval from 1,000 bootstrap replicates that resample whole clades (lineage prefix of depth seven in bacteria and eight in eukaryotes). Dotted vertical lines mark the proteome sizes at which fitted counts are compared (6,000 proteins in A and B, 20,000 in C and D). No phylogenetic regression was performed.

In plants the expansion largely tracks proteome size: the median plant proteome carries about 2.7-fold more SAM-methyltransferases than a non-plant eukaryote (371 versus 139), but it also has about 2.4-fold more proteins (34,300 versus 14,013), so SAM-methyltransferases make up a similar share of the proteome (median 1.14 versus 1.02 percent). In the clade-wise fits, the plant exponent is 0.69 [0.44, 0.80] for the total count and 0.75 [0.66, 1.54] for the Metabolite count (Figure 5C,D). At 20,000 proteins the fitted total count is 259 [236, 281] in plants, 236 [138, 272] in fungi, 192 [131, 252] in the remaining eukaryotes, and about 110 in metazoa. These data do not show that plants specialize in small-molecule methylation beyond what their proteome size predicts, because the fitted Metabolite count of fungi at comparable proteome sizes is similar to that of plants. Plants therefore carry more small-molecule methyltransferases in absolute terms (median 146 versus 47; Mann-Whitney p = 1.1e-21) mainly because the total methyltransferase count scales with proteome size, whereas the Metabolite fraction of the methyltransferasome increases only slightly (34 to 42 percent; p = 8.3e-3).

Actinomycetota, by contrast, devote most of a smaller methyltransferasome to small molecules: about 72 percent of a typical actinomycete’s SAM-methyltransferases are Metabolite, against about 35 percent for other bacteria (p = 7e-37), and the small-molecule count is 69 versus 17 (p = 1.3e-26). Both lineages have high small-molecule methyltransferase counts, but the expansion differs: in plants the total count scales with proteome size, whereas in Actinomycetota the Metabolite fraction of the methyltransferasome is also increased (72 versus 35 percent in other bacteria).

Within Actinomycetota the expansion is order-specific and tracks biosynthetic machinery rather than phylum membership. The load is driven by the biosynthetically rich orders (Pseudonocardiales, the Streptomyces-related Kitasatosporales, and Mycobacteriales), whose large genomes devote a substantial fraction to natural-product pathways studded with SAM-dependent tailoring methyltransferases. The reduced-genome order Micrococcales, soil- and plant-associated Actinomycetota such as Microbacterium, Arthrobacter, and Clavibacter, is a clear exception: it carries roughly a third of the total SAM-methyltransferase load of the other actinomycetes (median 33 versus 109 enzymes per organism, Mann-Whitney p = 1.3e-14) and a much smaller small-molecule fraction (35 versus 74 percent, p = 6.9e-15). Critically, the deficit is not a uniform shrinkage: it is concentrated almost entirely in the Metabolite substrates class, which falls to about a sixth of the other-actinomycete level (median 13 versus 82 enzymes), while the housekeeping RNA-modification machinery is nearly preserved (16 versus 18). That is exactly the signature expected if what is lost is secondary-metabolism tailoring rather than core methylation, consistent with the far fewer biosynthetic gene clusters reported for these organisms.^78,79^ SMSTPred thus recovers a known genomic and ecological distinction, biosynthetically rich versus streamlined Actinomycetota, from substrate-type predictions alone, indicating that it tracks metabolic capacity rather than taxonomy.

Among plants, the green algae (Chlorophyta; unicellular Chlorellales, Mamiellales, Chlamydomonadales) have the smallest methyltransferasome, carrying roughly 45 percent of the SAM-methyltransferase load of vascular land plants (median 191 versus 415; Mann-Whitney p = 2.9e-9) but a similar Metabolite fraction (39 versus 42 percent; p = 0.17) (Figure 4). Despite the smaller absolute methyltransferasome, the green algae devote a significantly larger share of their small proteome to SAM-MTases than other plants (median 1.49 versus 1.00 % of the proteome; Mann-Whitney p = 2.0e-9, Figure 3), therefore retaining much of the plant clade’s methylation machinery at higher density.

### The embedding space recovers known methyltransferase families

To see which proteins drive these trends, we projected all 115,850 methyltransferase embeddings into a shared UMAP, colored by their predicted substrate. The enzymes resolve into distinct, substrate-coherent islands (Figure 6), each an assembly of related enzymes. These islands are not arbitrary. Density-based clustering of the UMAP coordinates assigns each to a canonical family, reflecting similarity in the PLM embeddings.

**Figure 6.**
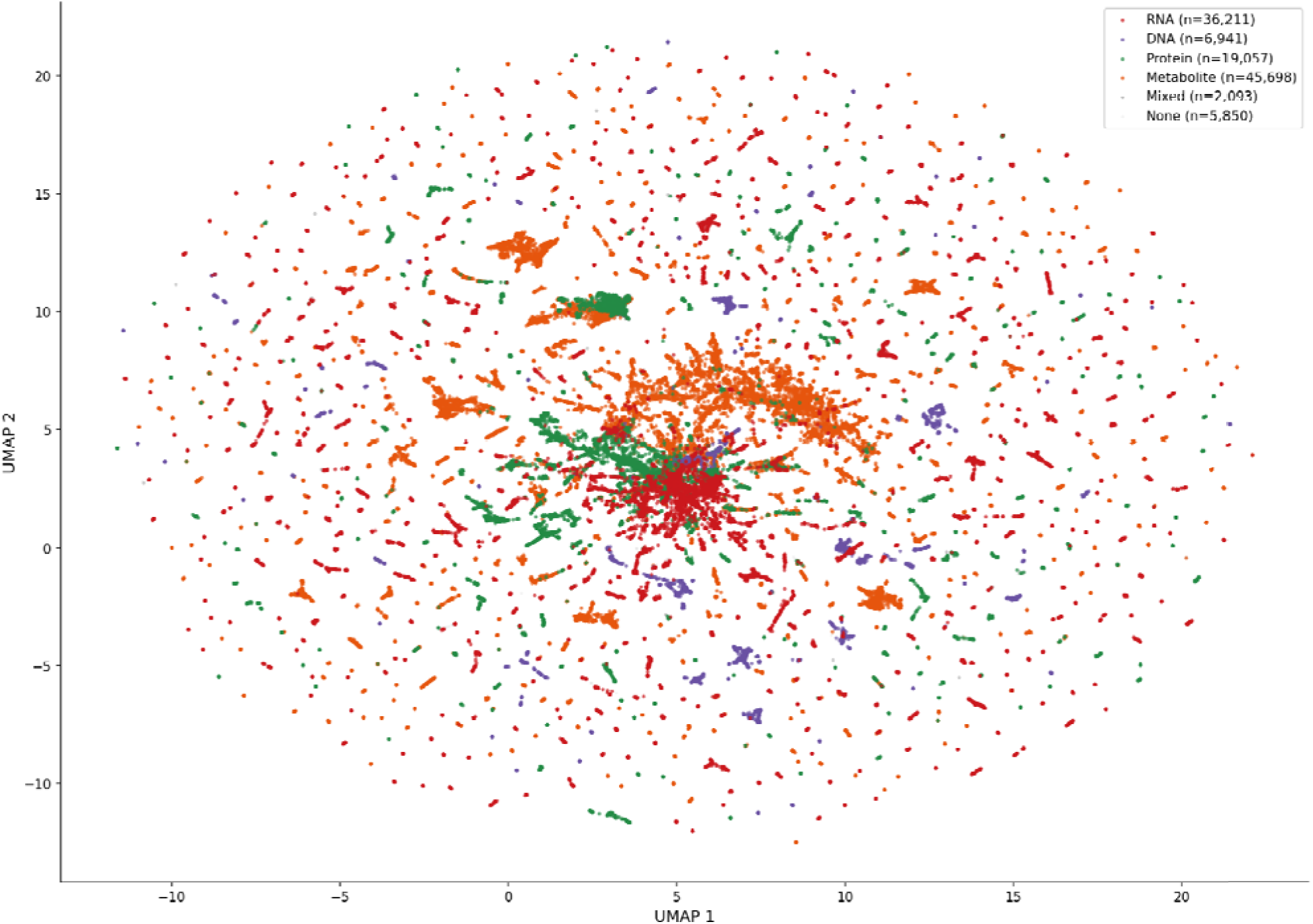
Substrate-colored global UMAP of 115,850 tree-of-life SAM-methyltransferases, which resolve into distinct, substrate-coherent islands; density-based clusters correspond to enzyme families (Figure 7). An interactive version is provided in the repository.

Zooming in on clusters enriched for predicted protein and DNA methyltransferases in Metazoa and Vertebrates, we see that most clusters are anchored by one or more reviewed (Swiss-Prot) members. Most of these reviewed members are PRMT and CARM1 arginine methyltransferases for Metazoa protein methyltransferases and DNMT1/3A/3B/3C cytosine methyltransferases for Vertebrate DNA methyltransferases (Figure 7A,B). This is evidence that SMSTPred is recovering consistent substrate specificity within families, and gives further evidence to the metazoan and vertebrate DNA and protein methylation trends.

**Figure 7.**
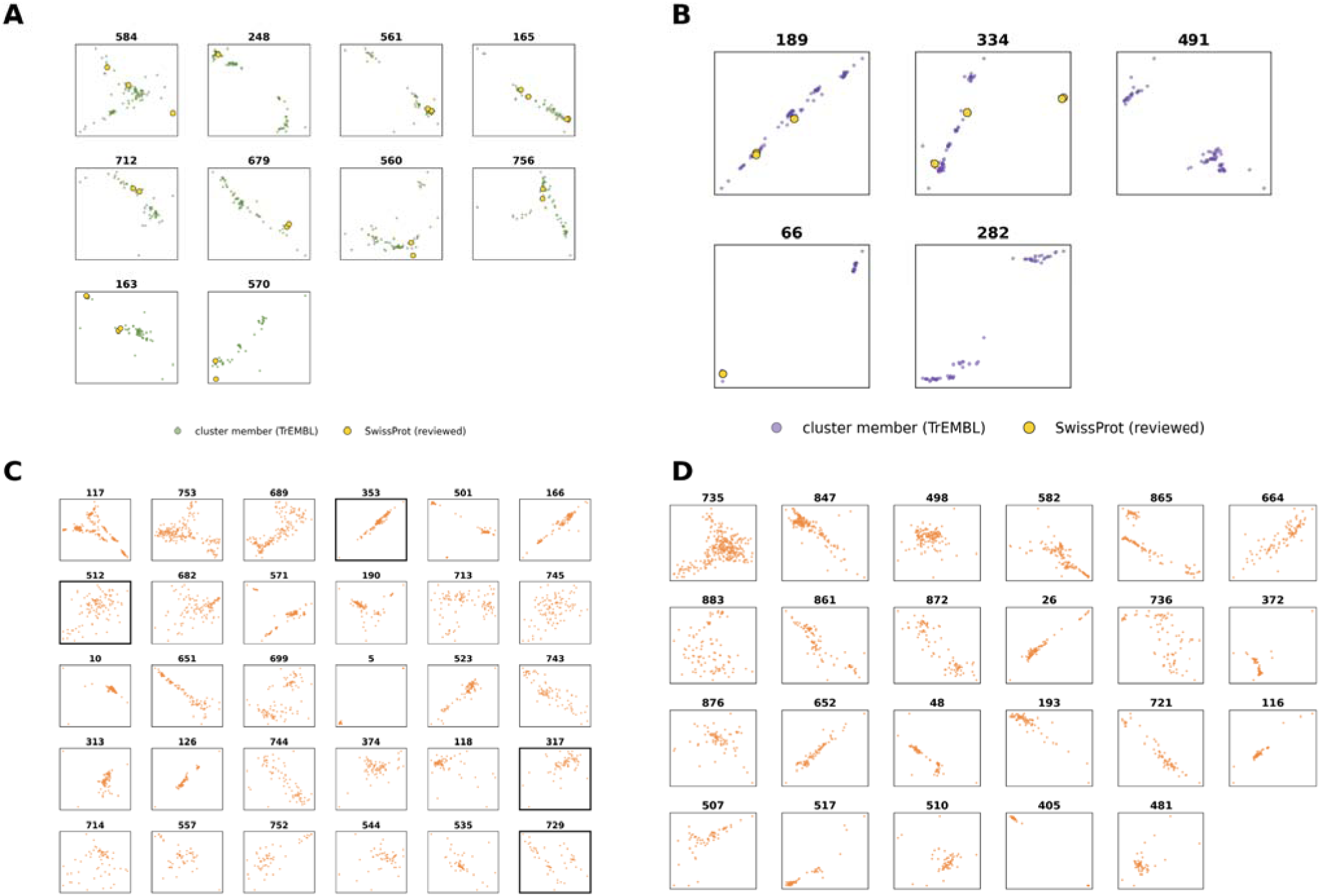
Family clusters underlying the trends, characterized and uncharacterized. Each panel is one HDBSCAN cluster of the ProtT5 UMAP of the tree-of-life methyltransferases, scaled to the extent of the cluster, with the cluster number above it. (A) Ten metazoan protein-methyltransferase clusters (PRMT5/7, HEMK2, ANT-KMT and others) and (B) five vertebrate DNA-methyltransferase clusters (DNMT1, 3A, 3B, 3C; two are unanchored), recovered de novo and anchored to reviewed Swiss-Prot members (yellow). (C, D) The 53 dark clusters, which contain no reviewed member and in which at least 80 percent of the proteins are from plants (C, 30 clusters) or Actinomycetota (D, 23 clusters) and predicted as Metabolite (small-molecule) methyltransferases (5,216 proteins in total). The four clusters examined in Figure 8 are outlined.

To compare the predictions to current UniProt annotations, the InterPro annotations were retrieved for each enzyme in the prediction set. Then, the in-class vs out-of-class fold-enrichment of each InterPro annotation was calculated for each predicted substrate class. Predicted substrate classes were cleanly enriched with known InterPro annotations that fit within each class: DNMT and C5-cytosine / restriction methylase domains for DNA; SET, PRMT, and HemK for Protein; ribosomal- and transfer-RNA modification domains for RNA; and O-methyltransferase and SABATH domains for Metabolite substrates (Table S3). This orthogonal, annotation-based validation supports the view that the predicted classes correspond to real biology.

### Substrate classification of uncharacterized MTases

Beyond the recovered families, the predictions assign a single substrate class to thousands of entirely unannotated enzymes, organized into discrete families with no reviewed (Swiss-Prot) members (Figure 7C,D). Only about half of these proteins (51 percent of 5,216) lie inside SMSTPred’s applicability domain (above 0.81 cosine similarity to the training proteins; see Methods), so the dark-cluster calls are best read as leads. The largest of these dark-matter groups are lineage-enriched small-molecule (’Metabolite substrates’) families in plants and actinomycetes, prime candidates for functional discovery and natural-product biosynthesis.^80^

We first assigned all 5,216 dark-cluster sequences a nearest-known-enzyme call with CLEAN (Supplementary Tables S1 and S2).^37^ To cross-validate these EC calls with a reaction-level signal, we separately screened the same dark-cluster sequences against all Rhea reactions using Horizyn v1.0. As a validation of the approach itself: with no hint that these are SAM-methyltransferases, Horizyn ranked a known SAM-methylation reaction as the single best match for 88.0 percent of proteins (in either reaction direction; 38.6 percent for the forward direction alone), and within the top 100 for 99.2 percent (5,172 of 5,216), with a median rank of 2 for each protein’s top-ranked SAM-methylation reaction. Because Rhea often encodes many near-duplicate reactions (for example, one reaction per isoprenoid chain length for menaquinone), we additionally clustered all reactions in Horizyn’s embedding space (UMAP + HDBSCAN, matching this study’s embedding-clustering convention) into reaction families, and report each dark cluster’s dominant family: every member’s individually best-scoring reaction, grouped by family, with the most common family’s share of the cluster and the fraction of members whose best-scoring reaction matched no coherent family (’noise’). Full results of both methods for all 53 clusters, the CLEAN call with its confidence tier and the Horizyn reaction family, are reported in Supplementary Tables S1 (Actinomycetota) and S2 (plants).

Four clusters stand out where CLEAN’s EC-centroid call and Horizyn’s dominant reaction family converge on a specific methylation chemistry (Figure 8). For c317 (n = 69), CLEAN assigns acetylserotonin O-methyltransferase at reliable confidence (d = 5.4), and 90 percent of members have the conversion of N-acetylserotonin to melatonin (RHEA:15573) as their own single best match. For c353 (n = 126), CLEAN’s reliable call (carnosine N-methyltransferase, d = 5.9) agrees with 80 percent of members ranking carnosine to anserine first (RHEA:14205). For c729 (n = 50), CLEAN assigns sterol 24-C-methyltransferase (d = 5.4) and 58 percent of members rank lanosterol to eburicol first (RHEA:52652). For c512 (n = 114), the two methods agree on the chemistry class but not on the specific reaction: CLEAN assigns desmethylxanthohumol O-methyltransferase (d = 5.8), whereas 80 percent of members rank the reticuline to laudanine O-methylation first (RHEA:10444); both are O-methylations of plant natural products. In each case two independent methods, one built on enzyme-sequence similarity and the other on reaction chemistry, name the same type of biochemistry for a cluster with no reviewed members.

**Figure 8.**
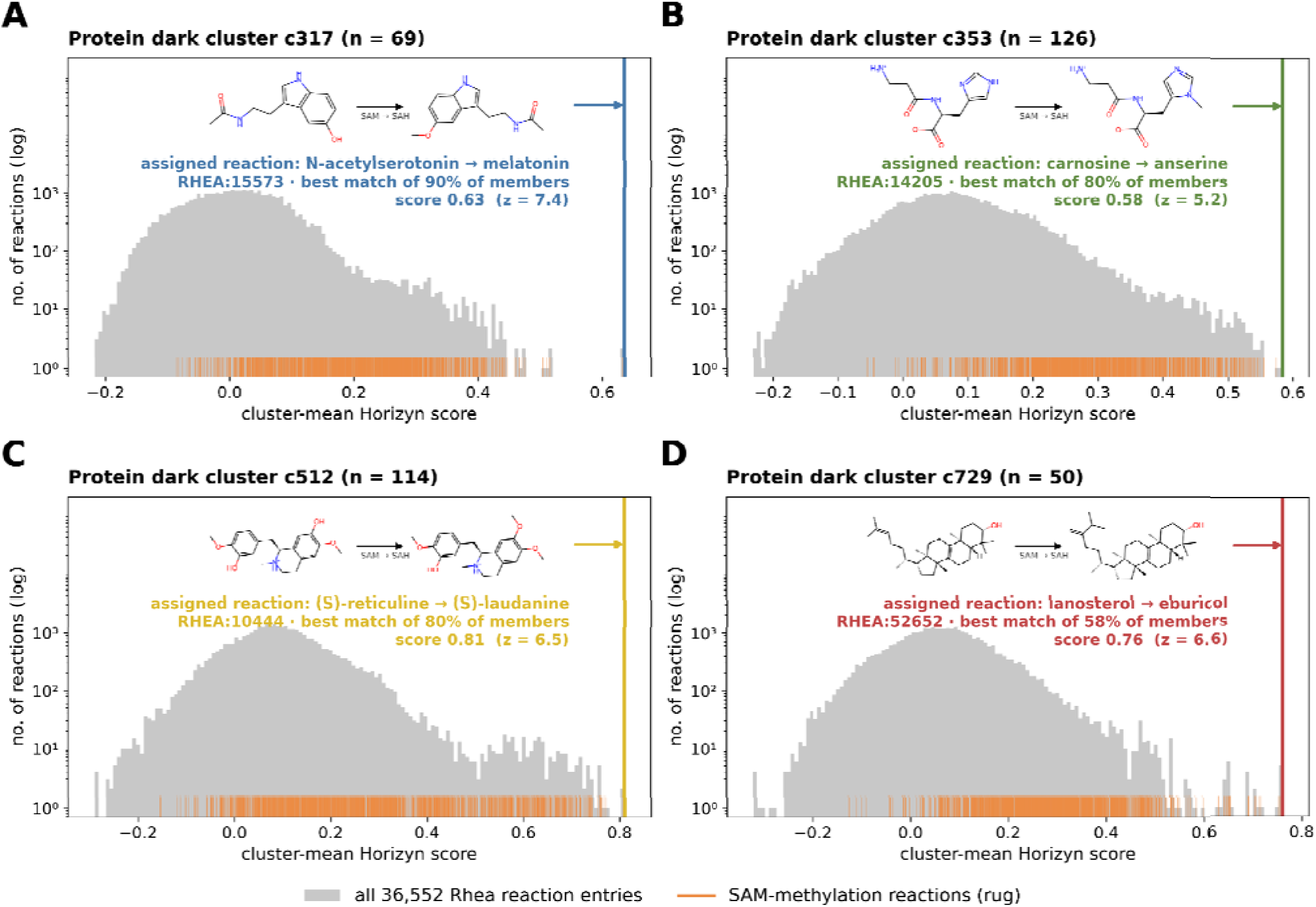
Horizyn cluster-mean scores of four plant dark clusters. Each panel shows, for one protein cluster (cluster index and number of proteins n), the distribution of the cluster-mean Horizyn score over all Rhea reaction entries (grey histogram, log axis), the SAM-methylation reactions (orange ticks), and the highest-scoring reaction (solid vertical line). The assigned reaction (substrate to product, SAM to SAH) is the best-scoring reaction of most members of the cluster; the panel gives its Rhea master ID, the percentage of members for which it is the best-scoring reaction, and the z-score of its cluster-mean score over all reactions. For comparison, the enzyme named by CLEAN for each cluster and the median distance d of its members to the EC centroid are given in the panel descriptions below; the CLEAN calls of all dark clusters are listed in Tables S1 and S2. (A), c317 (n = 69): N-acetylserotonin O-methylation to melatonin (RHEA:15573; 90% of members); CLEAN: acetylserotonin O-methyltransferase, d = 5.4. (B), c353 (n = 126): carnosine N-methylation to anserine (RHEA:14205; 80% of members); CLEAN: carnosine N-methyltransferase, d = 5.9. (C), c512 (n = 114): (S)-reticuline O-methylation to (S)-laudanine (RHEA:10444; 80% of members); CLEAN: desmethylxanthohumol O-methyltransferase, d = 5.8; the two methods name different specific reactions within the same O-methylation family. (D), c729 (n = 50): lanosterol C-24 methylation to eburicol (RHEA:52652; 58% of members); CLEAN: sterol 24-C-methyltransferase, d = 5.4. Reaction titles are generated from Rhea’s equations.

CLEAN also assigns nine dark clusters to enzymes that are not methyltransferases (clusters 664, 736 and 117 as alpha-onocerin synthase, 517 as spermidine synthase, 535 as spermine synthase, 166, 557 as nicotianamine synthase, 405 and 5; Supplementary Tables S1 and S2). For four of them (517, 535, 166 and 557) Horizyn independently agrees: all members have the same non-methylation reaction as their best-scoring reaction, for example the transfer of the aminopropyl group from decarboxylated SAM by spermidine and spermine synthase, or nicotianamine synthesis. These clusters entered the candidate set through InterPro SAM-binding domain membership (IPR029063), a criterion that also covers SAM-dependent enzymes that do not transfer methyl groups, such as the aminopropyl transferases; CLEAN, and for four of the clusters Horizyn, assign them non-methylation chemistry. The screen can therefore reject as well as support the annotation of a cluster as a methyltransferase.

### Sequence motifs explain only part of the substrate signal

For each class, STREME recovered the canonical catalytic and cofactor motifs: the Pro-Cys catalytic dyad for DNA (C5-cytosine chemistry) and the [D/N]PPY / NPPY tetrad shared by the DNA and Protein amino-methyltransferases, the PRMT ‘THW’ loop for Protein, and the m5C catalytic cysteine and a basic RNA-binding patch for RNA.

To test whether these motifs are class-discriminative rather than merely statistically enriched at discovery, we scanned all thirty-two against every one of the 122,807 proteins (FIMO, p < 1e-4) and computed each motif’s specificity as its occurrence rate in its own class divided by its occurrence rate in all other proteins pooled. Specificity ranged from about 13-fold (the DNA catalytic Pro-Cys) to near 1 (no enrichment), although every motif was significant at discovery. Twenty-four of the thirty-two are enriched at least two-fold over background (DNA 8, RNA 6, Protein 6, Metabolite 4), and were classified as class-discriminative. Their fold enrichment is given in Table 2. The remaining eight motifs are not class-discriminative: they are the pan-methyltransferase SAM-binding Motif I (the GxGxG glycine loop, recovered in both Metabolite and Protein at 1.3- and 1.4-fold) and several conserved Rossmann-core loops present in all classes. These motifs therefore reflect the shared SAM-binding machinery rather than the substrate and are excluded from the class-specific set.

**Table 1.** Classifier benchmark across all metrics (MCC, AUPRC, AUROC, F1, accuracy) for the nine representation and search-tool methods of Fig. 2A, cluster-disjoint split with the full training set, mean +/- SD over sixteen splits. Each metric is averaged over the four substrate labels. MCC, F1 and accuracy use a fixed threshold of 0.5, and AUPRC and AUROC are threshold-free. Rows are ordered by MCC. Embedding MLPs are shaded blue and search-tool baselines green, and the best value per metric is highlighted. The three best embedding MLPs (ProtT5, SaProt, ProtT5+ESM-IF) lie within 0.003 MCC of one another and do not differ significantly (Fig. 2A), so the per-metric highlights are not a ranking.

| Classifier | MCC | AUPRC | AUROC | F1 | Accuracy |
|---|---|---|---|---|---|
| MLP_ProtT5 | 0.940 ± 0.033 | 0.987 ± 0.013 | 0.997 ± 0.002 | 0.954 ± 0.028 | 0.986 ± 0.009 |
| MLP_SaProt | 0.938 ± 0.041 | 0.982 ± 0.017 | 0.996 ± 0.004 | 0.950 ± 0.035 | 0.986 ± 0.011 |
| MLP_ProtT5+ESM-IF | 0.937 ± 0.035 | 0.985 ± 0.016 | 0.996 ± 0.004 | 0.951 ± 0.029 | 0.985 ± 0.011 |
| MLP_ProstT5 | 0.895 ± 0.097 | 0.979 ± 0.018 | 0.995 ± 0.004 | 0.912 ± 0.082 | 0.980 ± 0.019 |
| Foldseek_1NN | 0.887 ± 0.044 | 0.925 ± 0.046 | 0.973 ± 0.015 | 0.913 ± 0.034 | 0.971 ± 0.018 |
| MLP_ProstT5+ESM-IF | 0.883 ± 0.085 | 0.967 ± 0.035 | 0.994 ± 0.004 | 0.899 ± 0.077 | 0.978 ± 0.018 |
| MMseqs2_1NN | 0.848 ± 0.058 | 0.925 ± 0.026 | 0.968 ± 0.015 | 0.881 ± 0.043 | 0.961 ± 0.021 |
| MLP_ESM-IF | 0.823 ± 0.089 | 0.919 ± 0.070 | 0.984 ± 0.013 | 0.854 ± 0.083 | 0.965 ± 0.017 |
| Profile_HMM | 0.519 ± 0.091 | 0.753 ± 0.061 | 0.781 ± 0.032 | 0.588 ± 0.079 | 0.803 ± 0.029 |

**Table 2.**
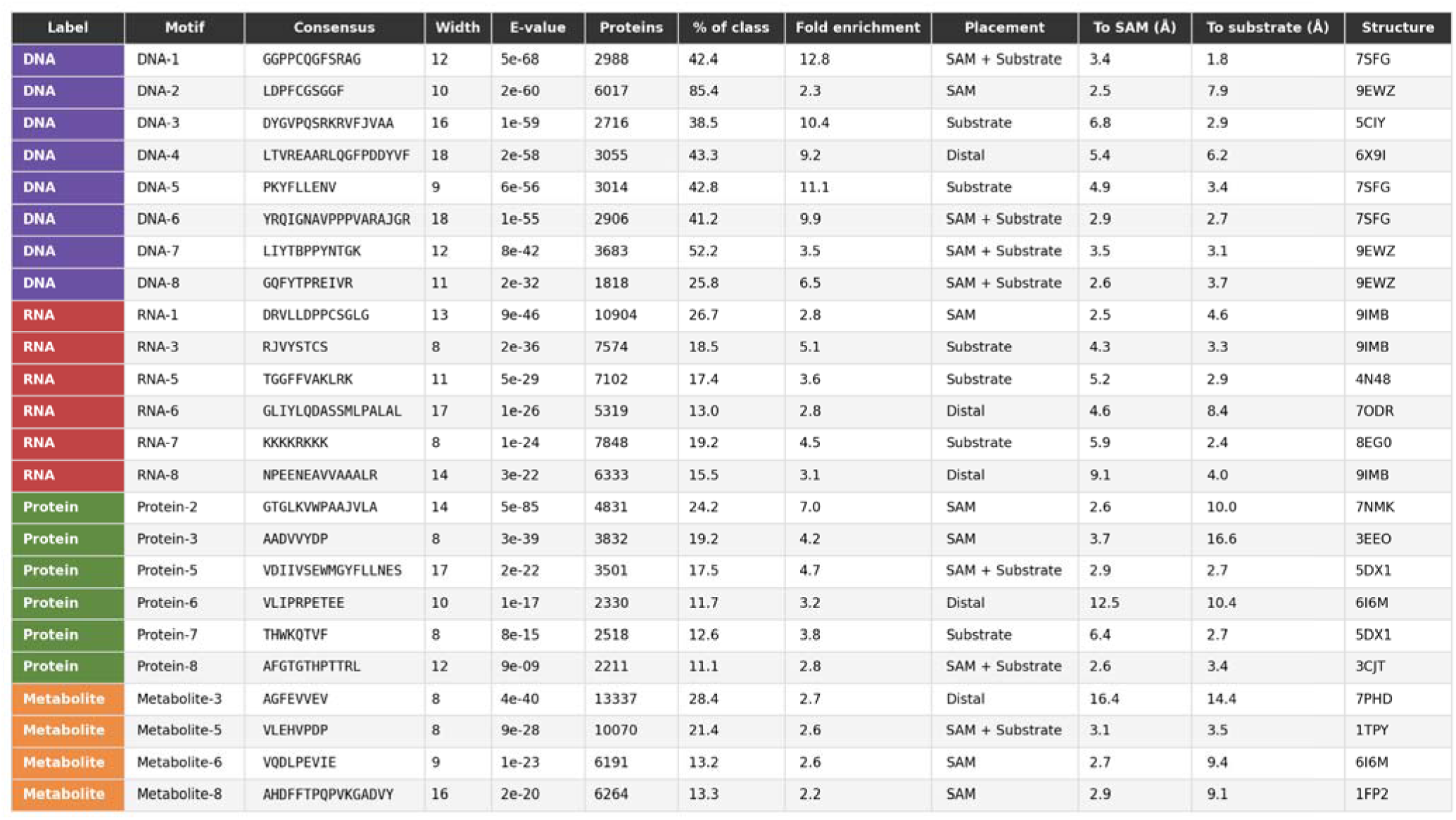
The twenty-four class-specific STREME motifs (of thirty-two discovered): consensus, motif width, discovery E-value, real per-protein coverage (number and percent of class members carrying each motif, from FIMO over all members), class specificity (occurrence rate in the motif’s own class divided by the occurrence rate in all other proteins pooled, over all 122,807 proteins), and structural placement (motif distance to the SAM cofactor and to the substrate in the selected structure; near substrate/SAM at 4 Å or less, distal beyond 4 Å from both). The eight pan-methyltransferase motifs below the specificity threshold are omitted.

To place the twenty-four class-specific motifs structurally, we selected for each, from the FIMO evidence, an experimental structure of a FIMO carrier that holds both a SAM/SAH cofactor and a substrate, located the motif by its matched sequence, and measured its minimum heavy-atom distance to the cofactor and to the substrate (Table 2). The majority are active-site elements: nineteen of the twenty-four lie within 4 Å of the substrate, the cofactor, or both (seven near both, six near the cofactor only, six near the substrate only), falling in catalytic and/or SAM-binding regions. Examples are the C5-cytosine catalytic loops of human DNMT1 (DNA-1 and DNA-6; Figure 9A,B), the PRMT substrate-binding loop of CARM1 (Protein-5; Figure 9C), and an active-site loop of the mycobacterial methyltransferase MmaA2 (Metabolite-5; Figure 9D).

**Figure 9.**
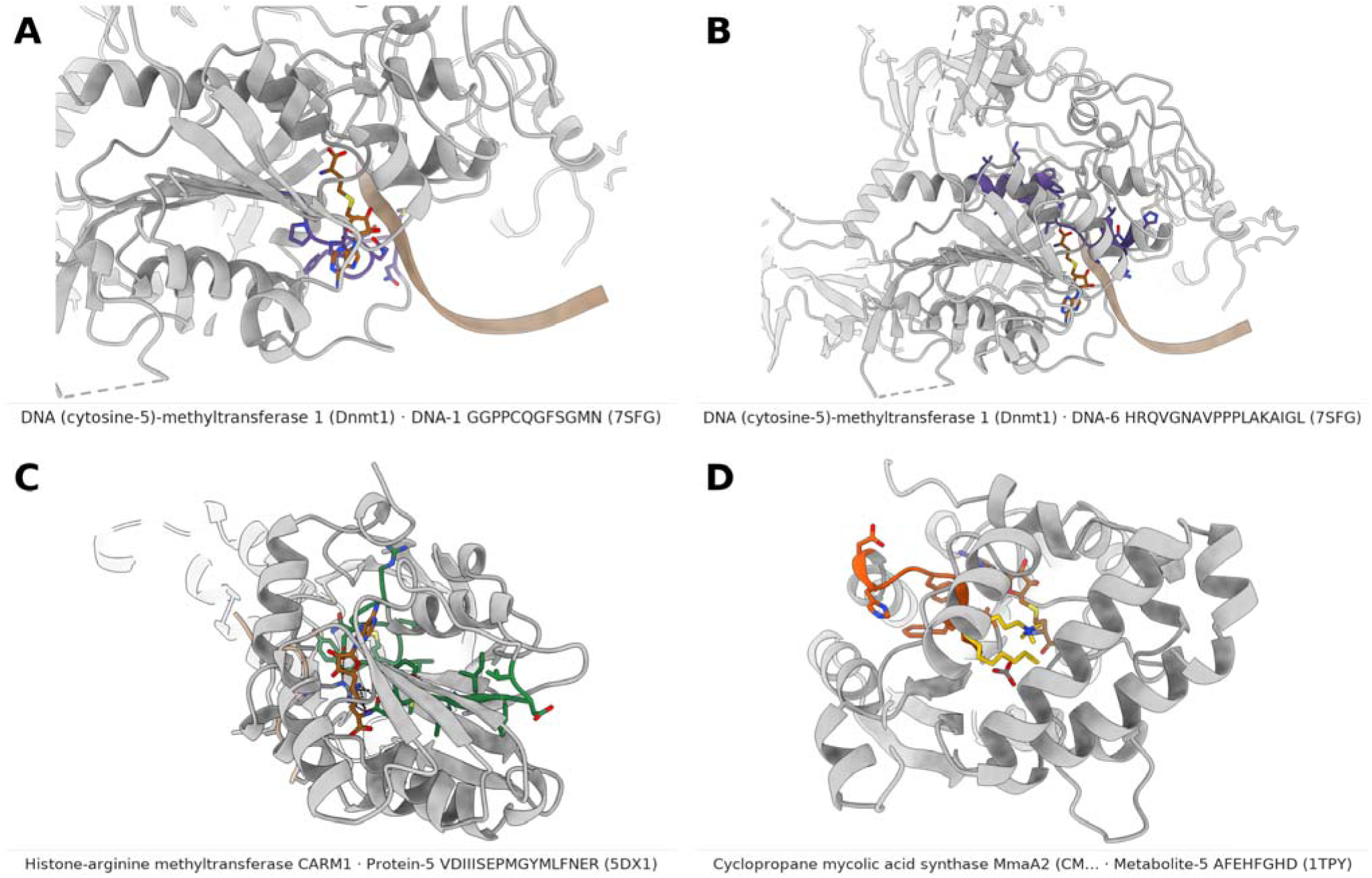
Class-specific motifs map onto characterized active sites. Each motif is shown on an experimental structure of an enzyme that carries it, with both the SAM/SAH cofactor and the substrate bound (grey enzyme, class-colored motif): (A) human DNMT1 (PDB 7SFG) with motif DNA-1 and (B) the same structure with motif DNA-6, (C) CARM1 (PDB 5DX1) with the PRMT substrate-binding loop (Protein-5), and (D) the mycobacterial methyltransferase MmaA2 (PDB 1TPY) with motif Metabolite-5. The structures were chosen by rule (see Methods).

Five class-specific motifs sit more than 4 Å from both ligands, four of which are shown in Figure 10: DNA-4 of DNMT1, RNA-6 of NSUN4, RNA-8 of NSUN6, and Metabolite-3 of the anthracycline methyltransferase DnrK. These motifs are distal only in the structure selected by our filter. In structures of other enzymes that carry the same motifs, they contact the substrate nucleic acid (RNA-8), partner proteins (RNA-6, in NSUN4 complexes, and Protein-6, in the PrmC complex with release factor 1) or the oligomer interface (DNA-4). Such motifs may therefore be conserved across enzymes while serving different functions in each, from substrate or partner recognition to assembly. This is inferred from deposited coordinates alone, and mutational analysis of these regions would be needed to test it.

**Figure 10.**
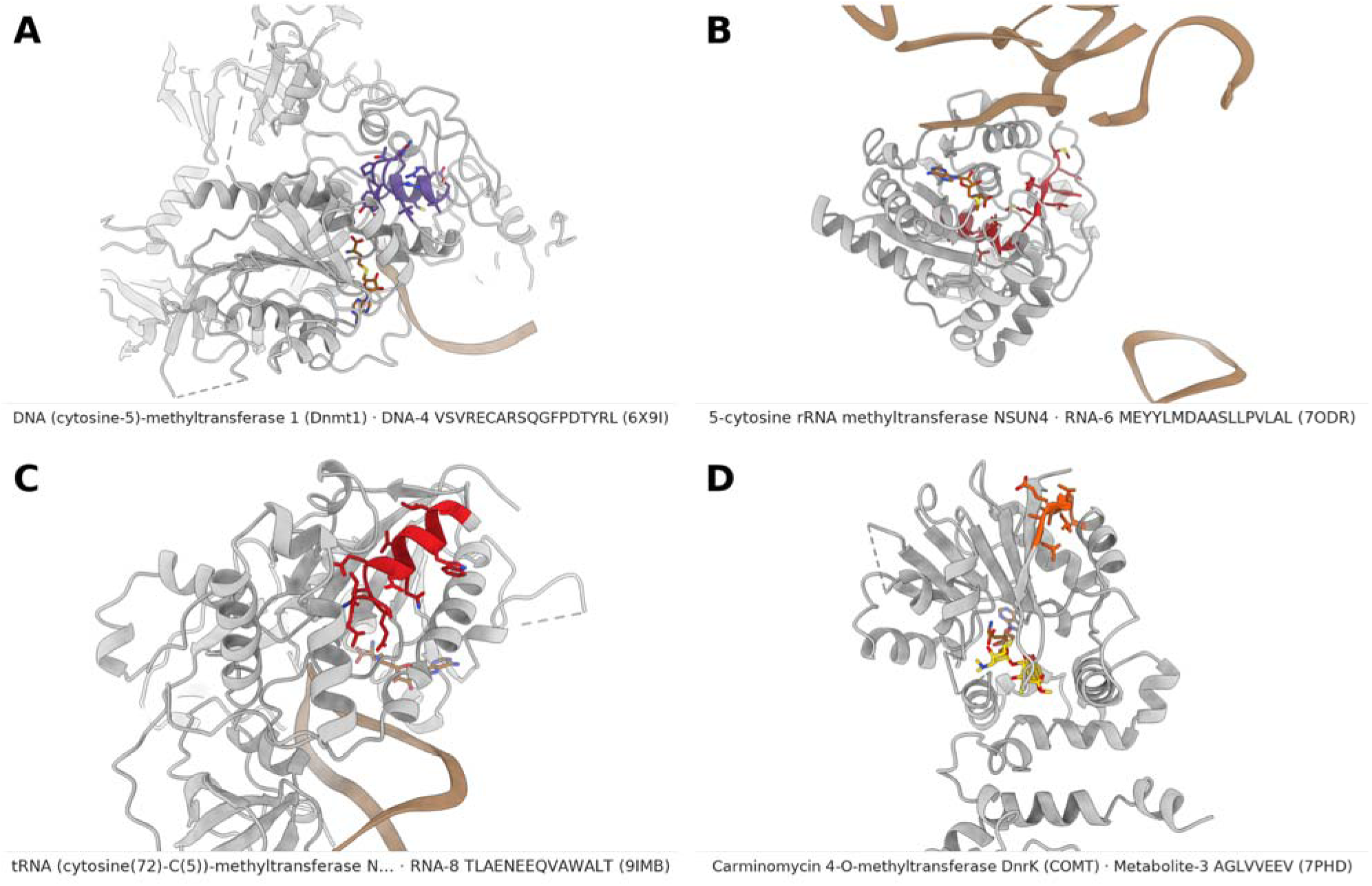
Class-specific motifs distal from both ligands (more than 4 Å from the cofactor and the substrate). (A) DNA-4 on DNMT1 (PDB 6X9I), (B) RNA-6 on NSUN4 (PDB 7ODR), (C) RNA-8 on NSUN6 (PDB 9IMB), and (D) Metabolite-3 on the methyltransferase DnrK (PDB 7PHD). These elements lie outside the catalytic and cofactor pocket and are candidates for substrate recognition or for an uncharacterized function.

Scanning every discovered motif over all 114,861 class members (FIMO, p < 1e-4), 11.1 percent of enzymes (12,732) match none of their class’s motifs despite carrying the SAM-methyltransferase fold; the deficit is concentrated in RNA enzymes (19.9 percent, versus 10.4 percent of Protein, 5.1 percent of Metabolite, and 1.4 percent of DNA). These motif-less enzymes are not a separate group: no embedding cluster is entirely motif-free, only seven clusters (675 proteins) are at least 90 percent motif-less, and motif-negative proteins lie predominantly within motif-bearing clusters (Figure 11), consistent with divergence within known families and not with distinct enzyme families. Nonetheless, SMSTPred recovers the RNA class under cluster-disjoint cross-validation (MCC 0.94), so for RNA enzymes substrate assignment must draw on sequence-derived features beyond the catalogued linear motifs.

**Figure 11.**
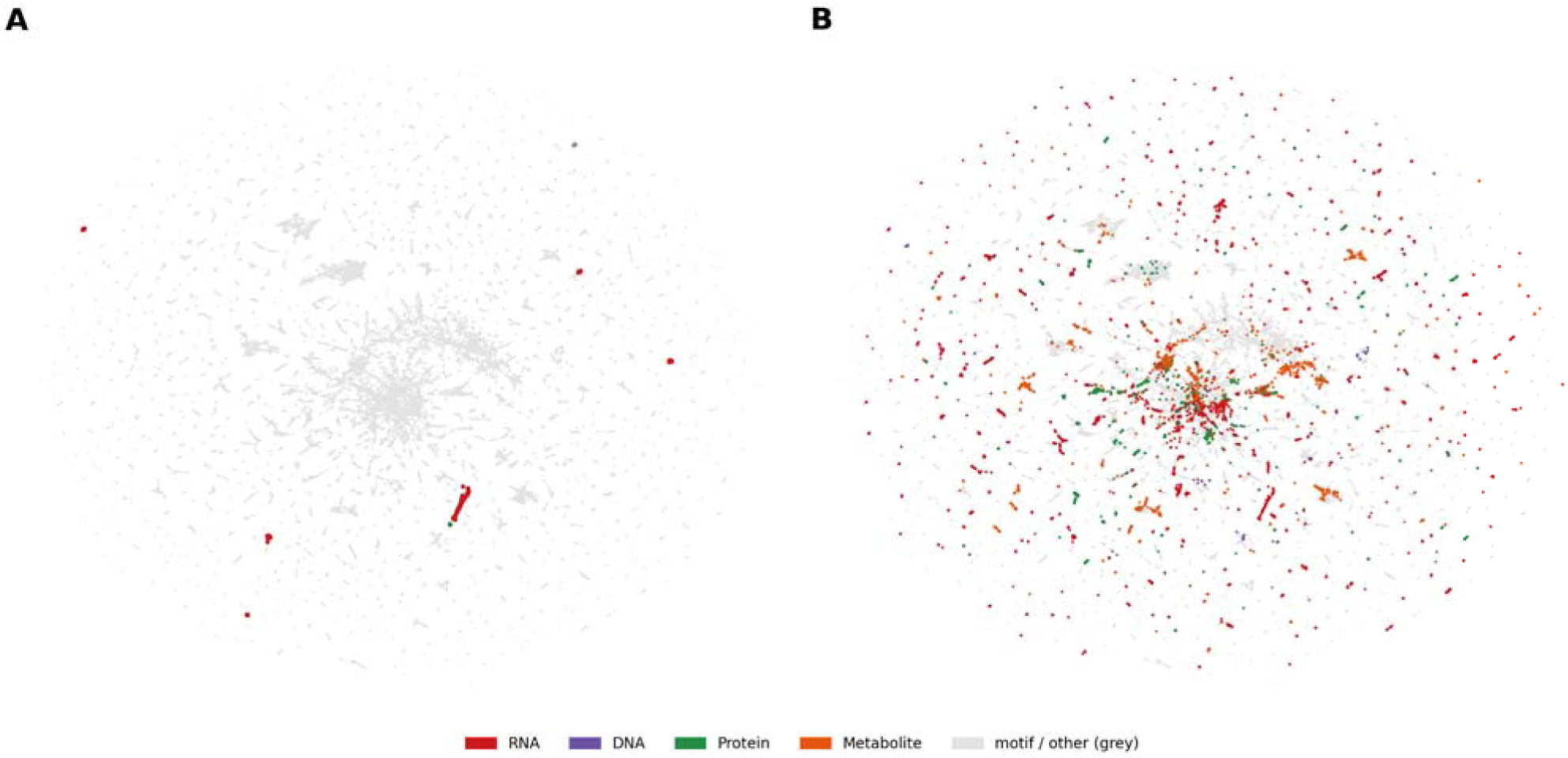
Enzymes carrying no discriminative motif (real per-protein FIMO). (A) The seven embedding clusters that are at least 90 percent motif-less (675 proteins); (B) all motif-less enzymes (11.1 percent of the 114,861 class members), colored by substrate, scattered within motif-bearing families rather than forming their own dark clusters.

## Discussion

SMSTPred predicts the substrate type of SAM-methyltransferases with high accuracy under homology-disjoint evaluation, and its application to 1,249 reference proteomes provides a resource of predicted substrate types. The lineage-level trends in these predictions are consistent with, but do not by themselves establish, differences in methylation strategy between lineages.

Embeddings capture recognition signals beyond detectable homology, and a plain sequence model is enough: ProtT5 injects no explicit fold information and needs no structure at inference, so it scales to whole proteomes, yet it matched the embeddings that require a structure (SaProt and the ESM-IF concatenation; 0.940 versus 0.938 and 0.937) and clearly beat both sequence-homology search (MMseqs2, 0.848) and structure-homology search (Foldseek, 0.887). The structure-only ESM-IF embedding was the weakest learned representation (0.823). Substrate type is therefore readable from sequence-level information without an explicit structure. Interpretability is consistent with this account: for most enzymes the class-specific motifs are active-site elements that recover the canonical catalytic and cofactor signatures (19 of 24 lie within 4 Å of a ligand), yet about 11 percent of enzymes carry none of them, especially within RNA methyltransferases, and are scattered within motif-bearing families. Therefore, the embedding could draw on family-specific signals beyond a short linear motif.

Substrate specialization is a readable evolutionary signal. The enrichment of small-molecule methyltransferases in two great chemical lineages, plants and actinomycetes, the vertebrate expansion of DNA methylation, and the broad conservation of RNA-modification machinery together sketch how methylation strategy tracks biology. The small-molecule expansion in Actinomycetota is not a result of proteome size: at equal proteome size they carry more Metabolite-substrate SAM-MTases than other bacteria, and their total count also rises more steeply with proteome size (Figure 5A,B). In plants the picture is different: the large methyltransferasome follows from large proteomes, since SAM-MTases make up a similar share of the proteome and the count scales with an exponent not distinguishable from that of other eukaryotes (Figure 5C,D); a higher level at equal proteome size is suggested but not resolved. In Actinomycetota, an expansion of the small-molecule methyltransferasome beyond what the proteome or the total methyltransferasome predicts indicates an expanded metabolism; in plants the small-molecule class is enriched in absolute terms and as a fraction of the methyltransferasome, but the expansion mostly accompanies larger proteomes. Together with the number of dark clusters enriched for these organisms’ Metabolite-substrate-methyltransferases, this highlights the importance of experimental validation of these metabolic pathways. Two groups are informative exceptions: the Micrococcales, which lose specifically the small-molecule class while keeping the RNA-modification machinery, and the green algae (Chlorophyta), which have a smaller methyltransferasome but a similar small-molecule fraction. SMSTPred recovering this known genomic and ecological distinction from substrate predictions alone, with the compositionally correct signature, is an internal validation that the predictions carry biological signals beyond phylogeny. The abundance of ‘Metabolite substrates’ methyltransferase predictions in the tree-of-life set likewise suggests that the RNA-biased composition of Swiss-Prot reflects curation history rather than natural abundance.

The number of genes in a functional category scales with genome size in a category-specific way, near-flat for housekeeping functions and steeply superlinear for regulatory ones, a regularity established across prokaryotes, extended to individual homology-defined families, and more recently surveyed across all three domains.^67,71–73^ Those studies count genes per functional category or per sequence family. A methyltransferase fold family, however, comprises enzymes of diverse substrate specificity, so a family-level count cannot resolve DNA-, RNA-, protein- and small-molecule-directed enzymes. Substrate-type prediction partitions one fold family by what its members act on, which makes it possible to ask whether these classes accumulate at different rates as proteomes grow. Our fits here compare one lineage with the rest of its domain rather than surveying exponents across categories; they show a steeper exponent only for the Actinomycetota total count, with the other differences in level (Figure 5).

By substrate-typing the unannotated majority, this prediction-annotated set prioritizes candidate natural-product biosynthesis enzymes and provides discrete, lineage-enriched dark-matter families as targets for experimental characterization. This resource contains more than we discuss here. We examine in detail only the few plant clusters whose substrates the two methods place with confidence, but there are 53 dark clusters (5,216 proteins) in plants and Actinomycetota alone. For all of them this resource provides substrate-class predictions and candidate reaction-level calls from CLEAN and Horizyn, and it provides a substrate-class prediction for every methyltransferase across the tree of life. These data could indicate novel RNA, DNA, Protein, or metabolic methyltransferase chemistry, and serve as a useful resource to guide experimentation on any family of SAM-methyltransferases.

A large share of the candidate methyltransferases (43.7 percent of the tree-of-life set) lies outside the applicability domain of SMSTPred, indicating how much of SAM-methyltransferase sequence space is sparsely represented in the labeled data and remains experimentally uncharacterized. The independent CLEAN and Horizyn analyses of the dark clusters point the same way: CLEAN’s call is reliable for only 10 of the 53 clusters and unreliable for 27, and the two methods converge on a specific chemistry for four clusters. The clusters with unreliable CLEAN calls are also those least covered by the training data (30 percent of their members lie inside the applicability domain, against 90 percent for clusters with reliable calls). The predictions therefore serve to prioritize orphan sequences and to propose candidate reactions, but assigning a reaction requires further experimentation.

In the reported run, 24 of the 32 STREME motifs are class-enriched (at least 2-fold); across STREME seeds and releases this number is 22 to 25, although which motifs are recovered varies more (12 to 21 of 32 recur between seeds). Many map onto active-site elements in experimental structures, which offers an interpretable view of conserved features that can be examined further. These motifs are, however, an indirect readout of the classifier: they were found in the sequences, not in the model, and about 11 percent of enzymes, 20 percent of the RNA methyltransferases, carry none of them. Motif-free RNA methyltransferases are nevertheless typed almost as accurately as motif-bearing ones (98.7 versus 99.7 percent out-of-fold accuracy), so the classifier appears to use sequence features beyond these motifs. Per-residue attribution of the ProtT5 embedding and sparse-autoencoder analysis of its representation would identify these features and test whether they coincide with the STREME motifs.

## Conclusion

SMSTPred predicts the substrate type of SAM-methyltransferases from protein language model embeddings with high accuracy under homology-disjoint evaluation. Applied to 1,249 reference proteomes, it gives lineage-level patterns of predicted substrate types and flags where the training data are sparse. The released classifier, with its applicability-domain flag, can be applied to newly sequenced genomes to annotate methyltransferase substrate types, to compare methylation profiles across lineages, and to prioritize uncharacterized enzymes, such as the dark clusters, for characterization in natural-product discovery.

## Author contributions

NR: conceptualization, data curation, formal analysis, investigation, methodology, software, validation, visualization, writing – original draft. DR: investigation, methodology, resources, writing – review & editing. RAM: methodology, software, validation. AS: data curation, investigation. JEH: formal analysis, methodology, software. AJ: conceptualization, funding acquisition, methodology, project administration, resources, supervision, writing – review & editing.

## Data availability

The data supporting this article (embeddings, label tables, structures, deployed model weights, tree-of-life predictions and analysis results; UniProt release 2026_03) are openly available at Zenodo, https://doi.org/10.5281/zenodo.23655998, under CC BY 4.0. The SMSTPred code is available at https://github.com/jinichlab/SMSTPred under the MIT licence and is archived at https://doi.org/10.5281/zenodo.24186207.

## Acknowledgements

The authors thank the HHMI Hanna Gray Fellowship (GT16787) and the UC San Diego NIH FIRST Grant (NCI 1UA54CA272220) for funding. We thank the UniProt, AlphaFold Protein Structure Database, Rhea and ChEBI teams for openly available data, and the developers of CLEAN and Horizyn for making their models available. AI disclosure: the AI coding assistant Claude Code (Anthropic) was used to assist with code development and testing, preparation of the code repository and data archives, checking of manuscript numbers against analysis outputs, and proofreading and copyediting of the manuscript text. All AI-generated code and text were reviewed, tested and verified by the authors, who take full responsibility for the content.

