## Supplementary Information for "SMSTPred: Protein Language Model Embeddings Predict SAM-Methyltransferase Substrate Specificity and Reveal Metabolite Methyltransferasome Enrichment in Plants and Actinomycetota"

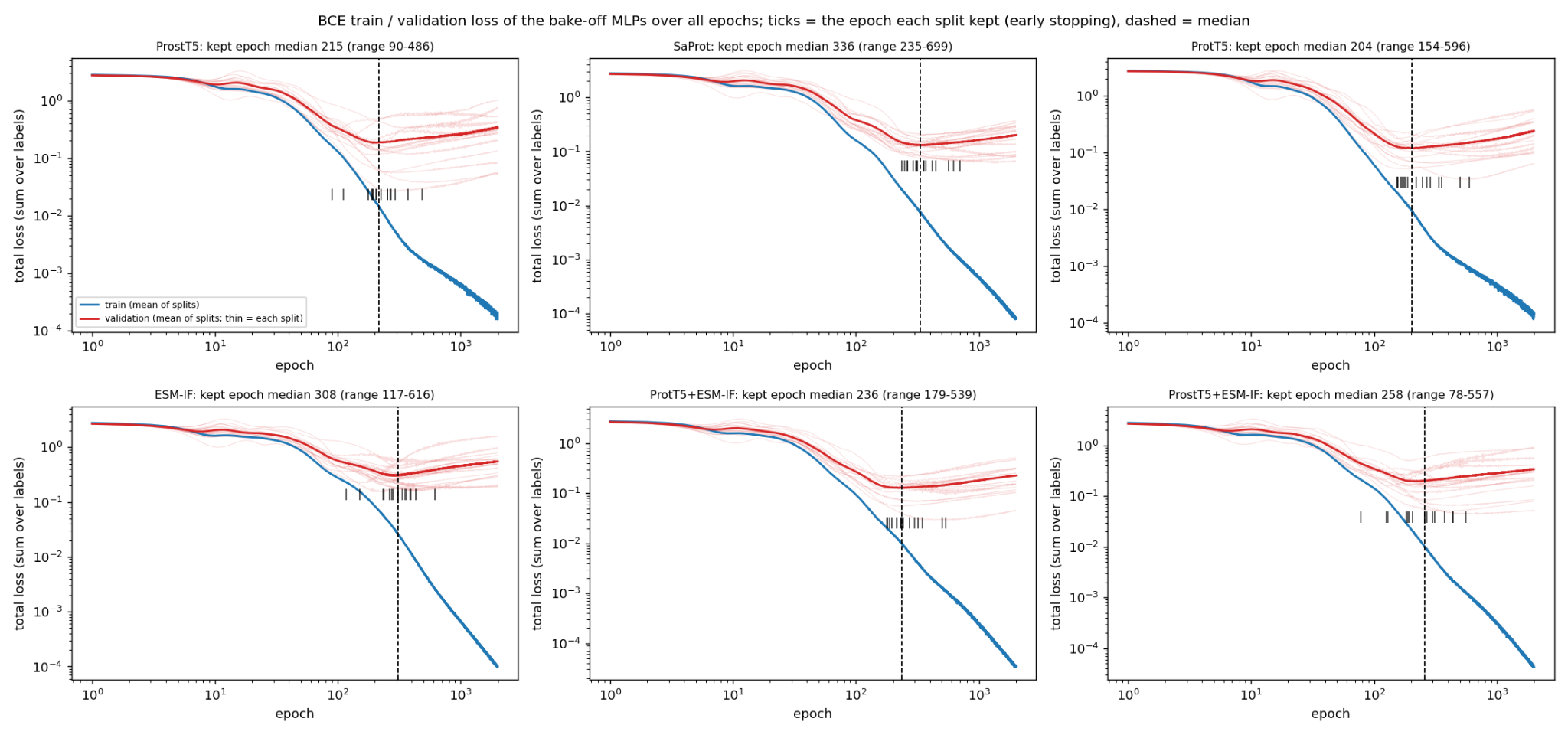

Figure S1. Training and validation loss of the six benchmark MLPs (ProstT5, SaProt, ProtT5, ESM-IF, ProtT5+ESM-IF, ProstT5+ESM-IF), trained on the MTase set. Mean binary cross-entropy loss per epoch on the training (blue) and validation (red; thin lines, each split) clusters over the sixteen cluster-disjoint splits. Ticks mark the epoch each split kept under early stopping (patience 80, at most 2,000 epochs) and the dashed line the median kept epoch (ProtT5 204, range 154 to 596).

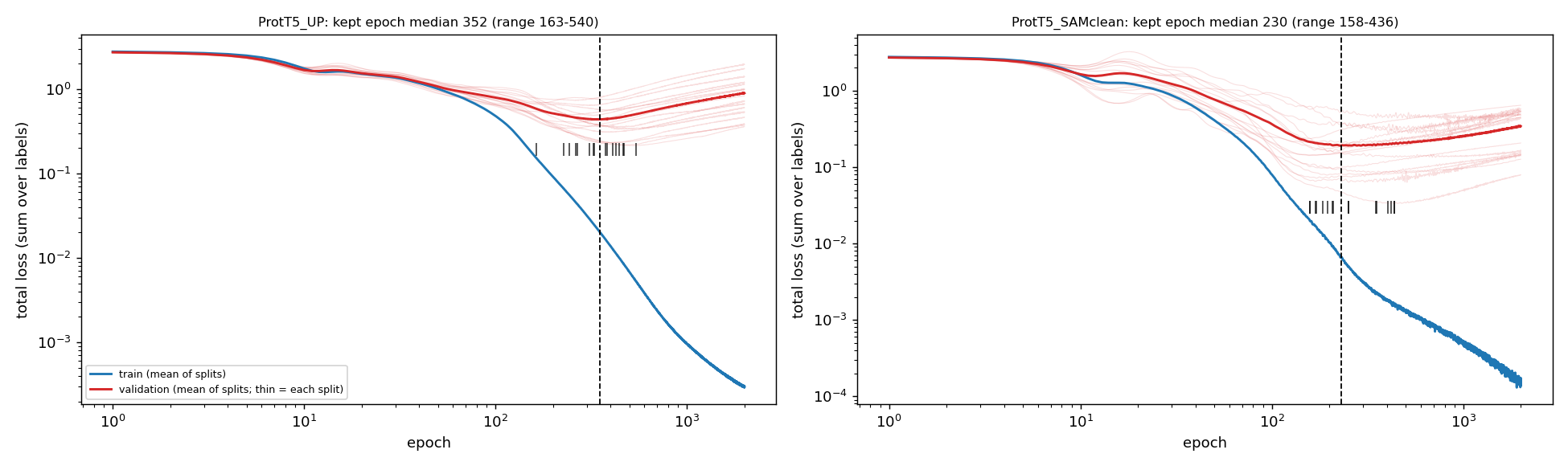

Figure S2. Training and validation loss of the UP-trained ProtT5 MLP (left) and the MTase-trained ProtT5 MLP (right; this sensitivity arm was retrained without the training and validation proteins that share a joint MTase and UP cluster with a test protein, leaving about 4,100 training proteins). Mean binary cross-entropy loss per epoch on the training (blue) and validation (red; thin lines, each split) clusters over the sixteen cluster-disjoint splits. Ticks mark the epoch each split kept under early stopping (patience 80) and the dashed line the median kept epoch (352 for UP-trained, range 163 to 540; 230 for MTase-trained, range 158 to 436).

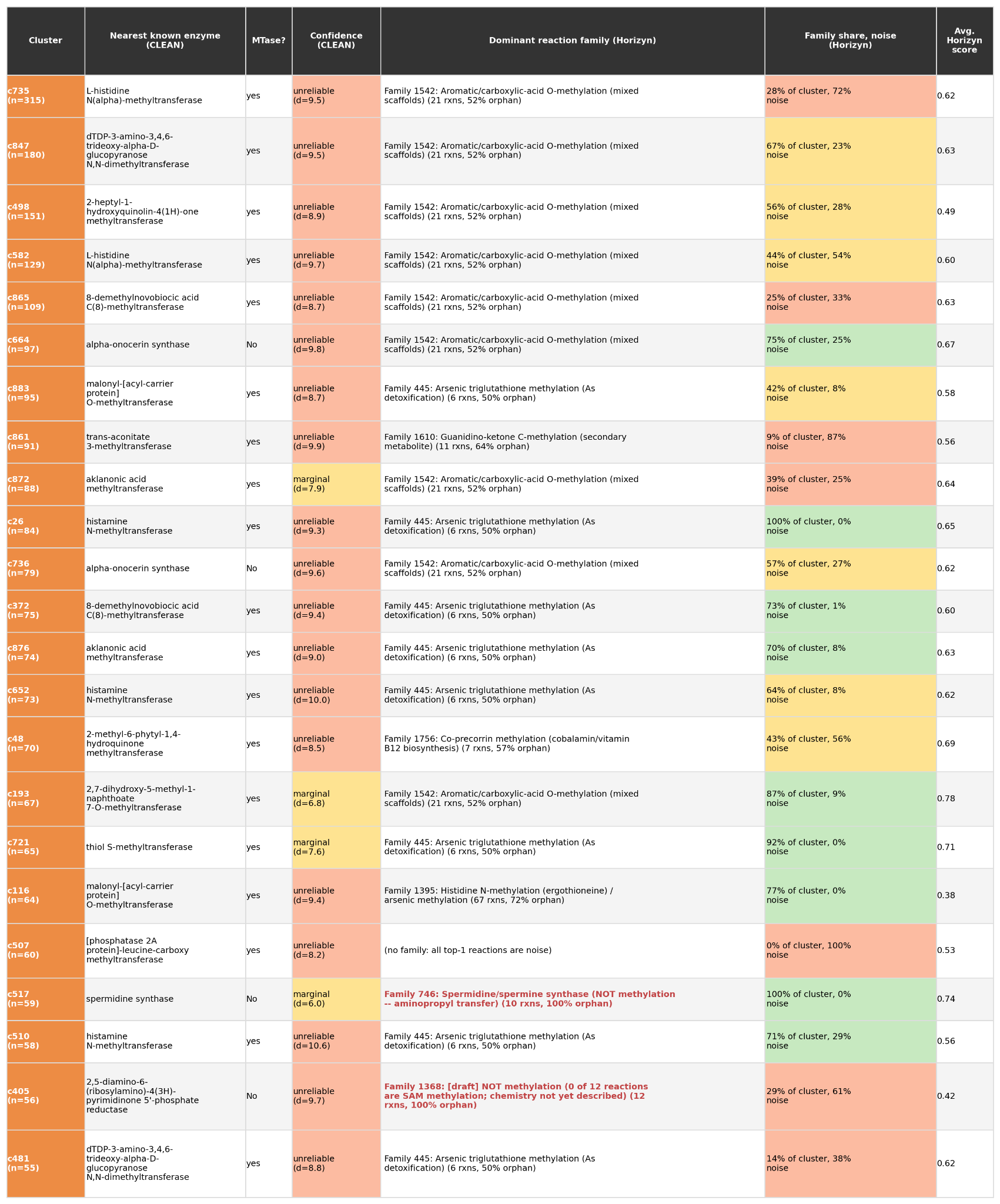

Table S1. Dark clusters of Actinomycetota (23 clusters; clusters with no reviewed member and at least 80 percent Actinomycetota proteins predicted as Metabolite methyltransferases). For each cluster: size (n), the nearest known enzyme from CLEAN with its call as a methyltransferase (MTase?) and confidence tier by median centroid distance d (reliable below 6, marginal 6 to 8, unreliable above 8), the dominant Horizyn reaction family with its size and fraction of orphan reactions, the share of the cluster in that family and the fraction of members whose best-scoring reaction matches no coherent family (noise), and the average Horizyn top-match score. Families flagged NOT methylation in red contain chemistry other than SAM methylation.

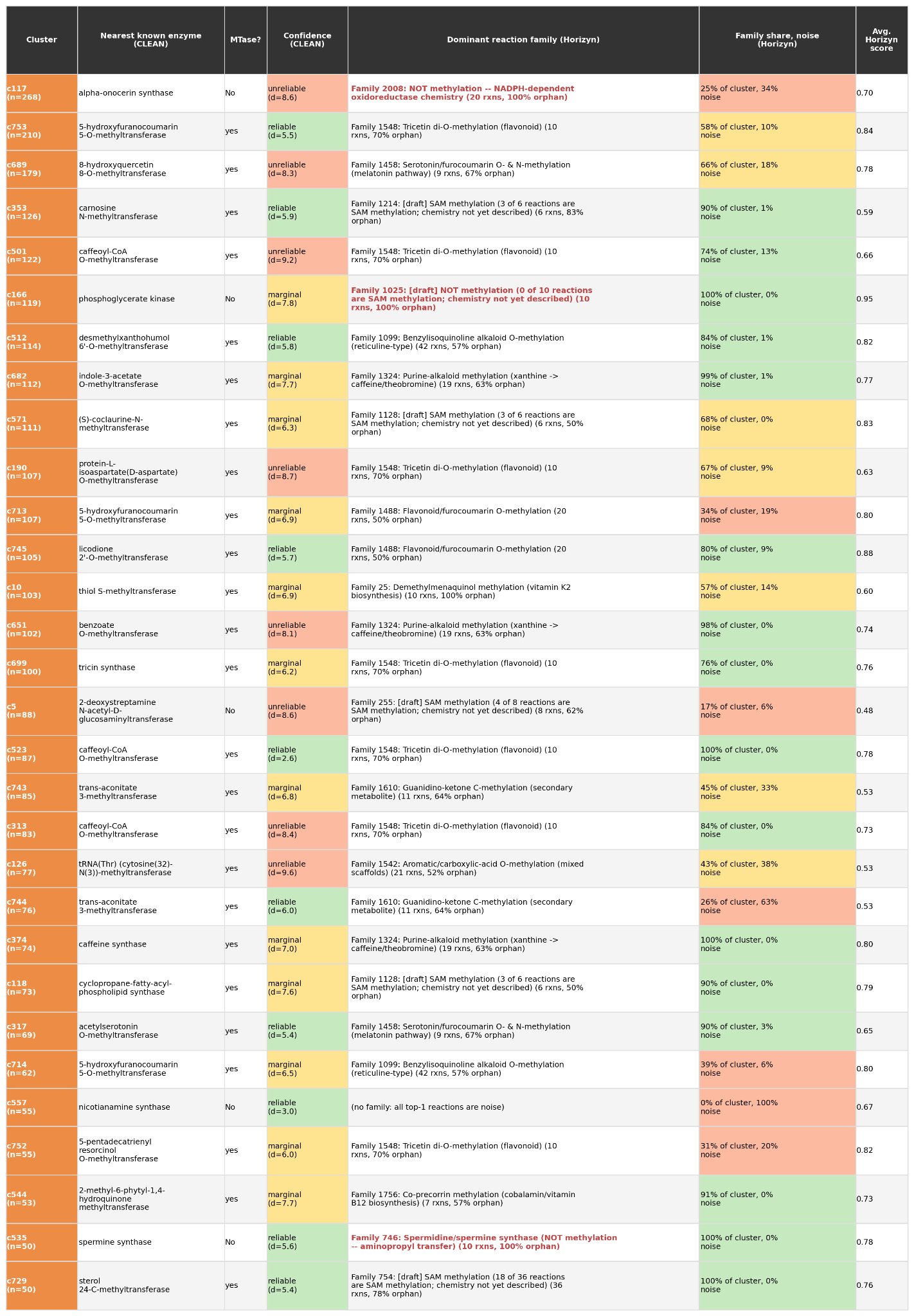

Table S2. Dark clusters of plants (30 clusters; clusters with no reviewed member and at least 80 percent plant proteins predicted as Metabolite methyltransferases), with the same columns as Table S1: size (n), the nearest known enzyme from CLEAN with its call as a methyltransferase (MTase?) and confidence tier by median centroid distance d (reliable below 6, marginal 6 to 8, unreliable above 8), the dominant Horizyn reaction family, the share of the cluster in that family and the fraction of members whose best-scoring reaction matches no coherent family (noise), and the average Horizyn top-match score. Families flagged NOT methylation in red contain chemistry other than SAM methylation.

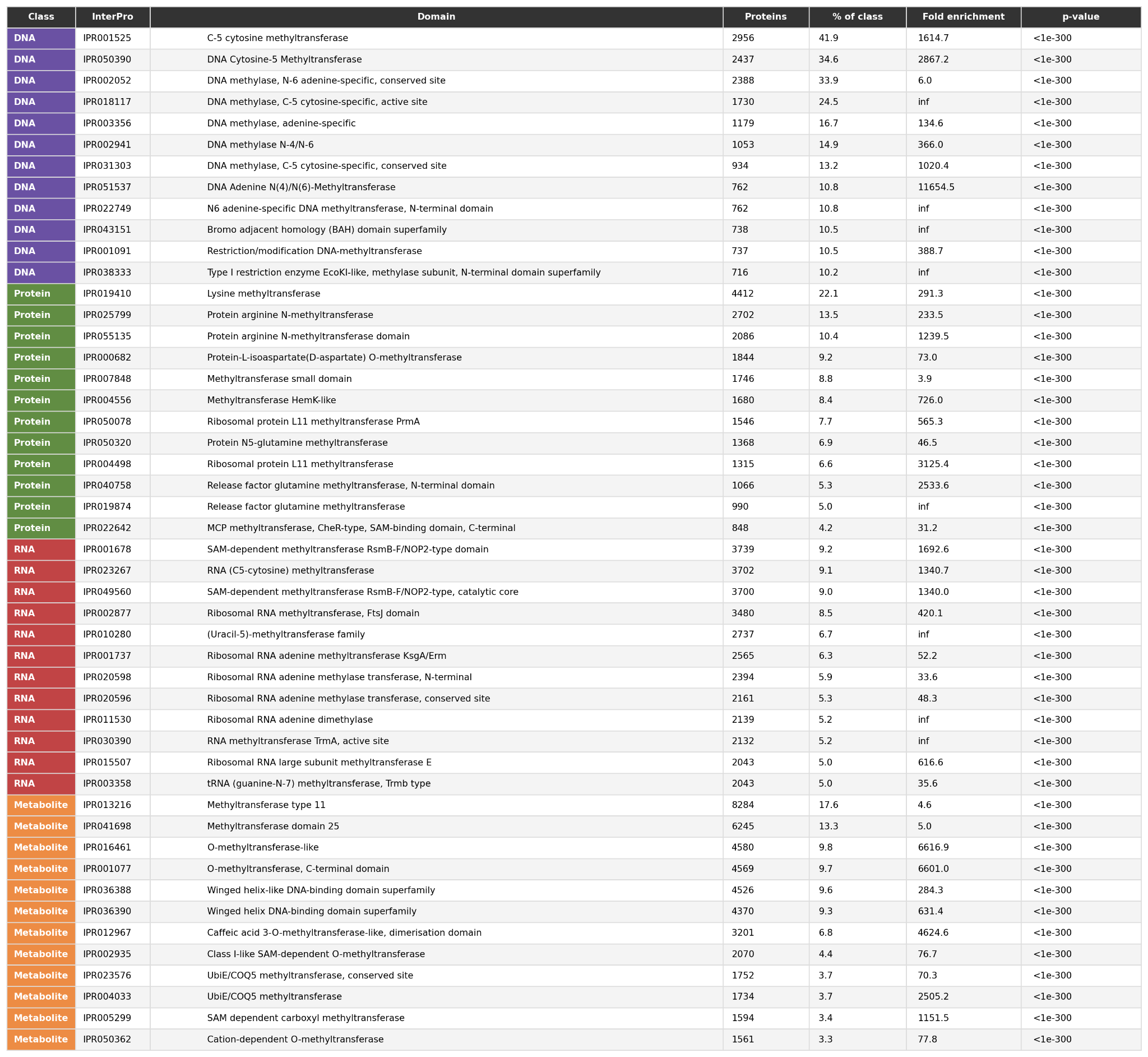

Table S3. InterPro domain enrichment of each predicted substrate class (DNA, Protein, RNA, Metabolite). For each class, the InterPro entries most characteristic of it: the entries with at least two-fold enrichment among that class’s members compared with all other classes, the twelve most common per class, without the pan-methyltransferase SAM-binding superfamily domain (IPR029063), which every class carries. Columns: the number of proteins of the class carrying the entry (Proteins), that number as a percent of the class (% of class), the fold enrichment (percent of the class over percent of the other classes; inf where no other class has the entry), and the p-value of a one-sided Fisher exact test of presence versus absence (below 1e-300 means smaller than the numeric range). Computed over the 114,861 proteins with a single class call.
